# Elevated CO_2_ alters soybean physiology and defense responses, and has disparate effects on susceptibility to diverse microbial pathogens

**DOI:** 10.1101/2024.06.04.595564

**Authors:** Melissa Bredow, Ekkachai Khwanbua, Aline Sartor Chicowski, Matthew W. Breitzman, Yunhui Qi, Katerina L. Holan, Peng Liu, Michelle A. Graham, Steven A. Whitham

## Abstract

- Increasing atmospheric CO_2_ levels have a variety of effects that can influence plant responses to microbial pathogens. However, these responses are varied, and it is challenging to predict how elevated CO_2_ (*e*CO_2_) will affect a particular plant-pathogen interaction. We investigated how *e*CO_2_ may influence disease development and responses to diverse pathogens in the major oilseed crop, soybean (*Glycine max* [L.] Merr.).
- Soybeans grown in ambient CO_2_ (*a*CO_2_, 419 parts per million (ppm)) or in *e*CO_2_ (550 ppm) were challenged with bacterial, viral, fungal, and oomycete pathogens, and disease, pathogen growth, gene expression and molecular plant defense responses were quantified.
- In *e*CO_2_, plants were less susceptible to *Pseudomonas syringae* pv. *glycinea* (*Psg*) but more susceptible to bean pod mottle virus, soybean mosaic virus, and *Fusarium virguliforme*. Susceptibility to *Pythium sylvaticum* was unchanged, although a greater loss in biomass occurred in *e*CO_2_. Reduced susceptibility to *Psg* was associated with enhanced defense responses. Increased susceptibility to the viruses was associated with reduced expression of antiviral defenses.
- This work provides a foundation for understanding of how future *e*CO_2_ levels may impact molecular responses to pathogen challenge in soybean and demonstrates that agents infecting both shoots and roots are of potential concern in future climatic conditions.

## Introduction

Atmospheric carbon dioxide (CO_2_) levels are steadily rising and by some estimates, are predicted to increase to 550 parts per million (ppm) or more by mid-century. Increasing CO_2_ levels, in combination with more extreme and unpredictable weather events, are expected to have significant impacts on global food security. The majority of plants on Earth use C3 photosynthesis, with relative atmospheric [O_2_] and [CO_2_] favoring photorespiration or photosynthesis, respectively (Pinto *et al*., 2014). As a result, elevated CO_2_ (*e*CO_2_) stimulates photosynthesis, generally increasing plant biomass and yield, providing a potential benefit to crop production (Dusenge *et al*., 2019; Ainsworth & Long, 2021). However, this CO_2_ “fertilization effect” depends on a number of factors, including water and nutrient availability and optimal growth temperatures (Ainsworth & Long, 2021). Moreover, suppression of photorespiration, once thought to be a wasteful process, has been associated with poor growth (Timm & Bauwe, 2013), lower nutrient and protein content (Taub *et al*., 2008; Broberg *et al*., 2017), and reduced abiotic stress tolerance (Voss *et al*., 2013) under ambient atmospheric conditions (*a*CO_2_).

Another possible impact of *e*CO_2_ is changes in the occurrence and severity of plant diseases. Pests and diseases that affect plant quality and yield pose one of the greatest challenges to crop production and are responsible for 17-30% of total yield losses of major crops (Savary *et al*., 2019). Climate change is expected to cause latitudinal shifts in disease pressure associated with the migration of important plant pathogens to new geographic regions (Bebber *et al*., 2013; Chakraborty, 2013; Raza and Bebber, 2022) as well as changes in pathogen virulence or aggressiveness (Pangga *et al*., 2007; Lake & Wade, 2009; Aguilar *et al*., 2015; Singh *et al*., 2023). Additionally, *e*CO_2_ has been demonstrated to alter interactions between plants and their attackers through changes in plant architecture, physiology, and molecular defense responses (Velásquez *et al*., 2018; Bazinet *et al*., 2022).

Plant disease development relies on three factors: a susceptible plant host, a virulent pathogen, and environmental conditions conducive to infection (Grulke, 2011). Over the past decade, tremendous advances have been made in our understanding of plant-microbe interactions (Ngou *et al*., 2022; Petre *et al*., 2022), however, it is still not well understood how changes in environmental conditions shape these interactions. Several mechanisms by which *e*CO_2_ affects defense responses have been reported, including changes in leaf nutrition (Ryalls *et al*., 2015; Sun *et al*., 2016), stomatal density (Li *et al*., 2015; Zhou *et al*., 2017), host metabolism (Matros *et al*., 2006; Ode *et al*., 2014), and redox homeostasis (Mhamdi & Noctor, 2016; Noctor & Mhamdi, 2017; Foyer & Noctor, 2020; Ahammed & Li, 2022). Altered levels of defense-related phytohormones, salicylic acid (SA), jasmonic acid (JA), and ethylene (ET), have also been reported (Zhang *et al*., 2015; Pan *et al*., 2019; Zhou *et al*., 2019). In general, constitutive and/or pathogen-induced SA levels are higher in the foliar tissues of C3 plants grown under *e*CO_2_ (Bazinet *et al*., 2022), suggesting increased resistance to viral and biotrophic pathogens (Vlot *et al*., 2009; Murphy *et al*., 2020). However, the extent and directionality of these responses are highly variable between studies (Bazinet *et al*., 2022), which may reflect species, cultivar or ecotype-specific CO_2_ adaptations, as well as differences in additional environmental growth conditions. For example, in tomato (*Solanum lycopersicum*), *e*CO_2_ increased SA biosynthesis, and plants displayed higher resistance to leaf curl virus, tobacco mosaic virus, and the hemibiotrophic bacterial pathogen *Pseudomonas syringae* pv. *tomato* (*Pto*DC3000), but were more susceptible to the necrotrophic fungal pathogen *Botrytis cinerea* (Huang *et al*., 2012; Zhang *et al*., 2015). In contrast, *e*CO_2_ increased JA-dependent responses in Arabidopsis (*Arabidopsis thaliana*), enhancing resistance to *B. cinerea* and reducing resistance to *Pto*DC3000 (Zhou *et al*., 2019). These discrepancies highlight the need for direct investigations of pathosystems involving important crop species rather than relying on generalized information from model systems.

Soybean (*Glycine max* [L.] Merr.) is the most widely grown protein and oilseed crop, accounting for approximately 329 million tons of global crop production in 2022 (http://www.worldagriculturalproduction.com/). The effect of *e*CO_2_ on soybean physiology has been investigated at various scales (Ainsworth & Long, 2021; Li *et al*., 2021). In general, soybean yield potential and root nodule mass increase under *e*CO_2_ (Dermody *et al*., 2008) with no significant impact on plant nutrient or protein content (Myers *et al*., 2014). *e*CO_2_ has also been associated with changes in leaf endophyte communities (Gonçalves *et al*., 2021; Christian *et al*., 2021), rhizospheric bacterial communities (Yu *et al*., 2016), and responses to herbivore attack (Casteel *et al*., 2008; O’Neill *et al*., 2010; Paulo *et al*., 2020). However, only one study has reported the effect of predicted future atmospheric conditions on soybean diseases by monitoring the incidence and severity of some naturally occurring diseases at the Soybean Free Air Concentration Enrichment (SoyFACE) facility (Eastburn *et al*., 2010; Aspray *et al*., 2023). As soybean is susceptible to numerous economically important pathogens, including bacteria, fungi, oomycetes, viruses, and nematodes (Whitham *et al*., 2016), a formal investigation of how *e*CO_2_ impacts susceptibility to diverse pathogens is needed. Moreover, the effect of *e*CO_2_ on molecular defense responses has not yet been investigated in this species.

In the present study, we compared the effects of CO_2_ levels on soybean defense response in plants grown under the current [*a*CO_2_] of 419 parts per million (ppm) versus an [*e*CO_2_] of 550 ppm. Using the soybean-*P. syringae* pathosystem, we assessed innate immune responses and disease susceptibility and conducted transcriptomic analyses to gain a global understanding of how *a*CO_2_ levels affect interactions in this pathosystem. We also conducted infection experiments using two foliar viral pathogens, bean pod mottle virus (BPMV) and soybean mosaic virus (SMV), and two filamentous root pathogens, *Fusarium virguliforme* and *Pythium sylvaticum*, and demonstrate that *e*CO_2_ exerts differential effects on interactions with these diverse microbial pathogens. Our work provides a foundation for future studies investigating the molecular interplay that regulates defense responses to *e*CO_2_ and identifies pathogens of potential concern in predicted future atmospheric conditions.

## Materials and Methods

### Plant growth and maintenance

Soybean (*G. max* cv. Williams 82) plants were grown in controlled environment chambers at the Iowa State University Enviratron facility (Bao *et al*., 2019) with lighting, humidity, and temperature conditions as illustrated in Figure S1. [CO_2_] maintained at 419 ppm represented *a*CO_2_ and 550 ppm represented future atmospheric conditions that may be reached by 2050 (*e*CO_2_) (Jaggard *et al*., 2010). For each condition, three replicate growth chambers were used to provide biological replicates. Plants were grown in LC-1 potting soil mix (Sungro) and fertilized weekly with 15-5-15 Cal-Mag Special (Peters Excel, G99140).

### Physiological Measurements

To assess changes in soybean physiology associated with *e*CO_2_, shoot biomass, photosystem II (PSII) activity (quantum yield of fluorescence; ΦPSII activity) and stomatal conductance (gas exchange rate; g_sw_) were measured for 15 plants per CO_2_ treatment. ΦPSII and g_sw_ were obtained using a LI-600 Portable System (LI-COR Biosciences, Lincoln, NE, USA) during the VC-V4 growth stages (Kumudini, 2010). ΦPSII measurements were conducted on the adaxial surface of unifoliate leaves of 2-week-old plants and the newest fully expanded trifoliolate leaves of 3- and 5-week-old plants. g_sw_ measurements were simultaneously collected from the abaxial surface. At 21 days after planting (dap), leaves were collected from eight plants per treatment and chamber, yielding three biological replicates per treatment for QuantSeq analyses (48 samples for QuantSeq dataset 1). Samples were collected between 09:00 and 10:00 am for all replicates. At 35 dap, shoots were collected to determine fresh weight and dry weight.

### Stomatal density, aperture, and index measurements

Stomatal density and aperture measurements were assessed from 10 leaf samples, while stomatal index measurements were taken from five leaf samples for each CO_2_ treatment at 21 dap. Impressions were made from the abaxial leaf surfaces using clear nail varnish (Ceulemans *et al*., 1995). Stomata were imaged using light microscopy, and counted in three randomly selected fields of view per leaf sample. Stomatal aperture and stomatal index measurements were conducted as described previously (Zhu *et al*., 2021; Sultana *et al*., 2021).

### RT-PCR and RT-qPCR analysis

Total RNA was isolated from approximately 50 mg of leaf tissue using TRIzol Reagent (Thermo Fisher Scientific, Waltham, MA, USA). Subsequently, 2 µg of RNA was reverse transcribed using the Maxima First Strand cDNA Synthesis Kit (Thermo Fisher Scientific, Waltham, MA, USA). Real-time quantitative PCR (RT-qPCR) was conducted using 2X PrimeTime Gene Expression Master Mix (Integrated DNA Technologies, Coralville, IA, USA) and multiplexed probes designed for *Pathogenesis-Related Protein 1* (*PR1*) (Glyma.13G251600), *Kunitz Trypsin Inhibitor 1* (*KTI1*) (Glyma.08G342000), BPMV, or SMV (Table S1). Expression of *Dicer-Like 2* (*DCL2*) (Glyma.09g025300) and *Argonaute 1* (*AGO1*) (Glyma.09g167100) was assessed using iTaq Universal SYBR Green Supermix (BioRad, Hercules, CA, USA). To quantify *F. virguliforme* and *P. sylvaticum*, soybean roots were collected at 14 and 35 dap, respectively. DNA was extracted using the CTAB method (Rogers & Bendich, 1994), and RT-qPCR was performed using primers specific to each pathogen (Table S1). *S-phase kinase-associated protein 1* (*Skp1*) (Glyma.08G211200) was used as an internal reference for all experiments (Beyer *et al*., 2021). Primers and probes were designed from cultivar Williams 82 reference sequence (version Wm82.a4.v1, (Schmutz *et al*., 2010)) using PrimerQuest Tool (Integrated DNA Technologies).

### Immune signaling assays

Oxidative species production was monitored in two-week-old soybean plants, as described previously (Bredow *et al*., 2019) using leaf discs collected from 16 plants per CO_2_ treatment. The elicitor solution contained 100 µM luminol (Sigma-Aldrich, St. Louis, MO), 10 µg/mL horseradish peroxidase (Sigma-Aldrich, St. Louis, MO), with or without 100 nM flg22 (VWR International, Radnor, PA). Chemiluminescence was quantified on a GloMax^Ⓡ^ plate reader (luminescence module; Promega, Madison, WI) every 2 min for 30 min with a 1000 ms integration time.

Flg22-induced MITOGEN-ACTIVATED PROTEIN KINASE (MAPK) activation was assessed as described previously (Xu *et al*., 2018) using leaf discs collected from six plants per CO_2_ treatment. Leaf discs were maintained in CO_2_ chambers, treated with 100 µM of flg22 peptide and sampled between 0 and 250 min. Total protein was extracted and normalized as described previously (Wang *et al*., 2018) and MAPK activation was assessed by immunoblot analysis using primary anti-Phospho-p44/42 MAPK antibody (Erk1/I 2; Thr-202/Tyr-204) (Cell Signaling, Danvers, MA, USA) and secondary goat anti-rabbit-HRP antibody (Cell Signaling, Danvers, MA, USA).

### Bacterial infection assays

*P. syringae* pv*. glycinea* (*Psg*), *Pto*DC3000, and *Pto*DC3000 *hrcC-* (*hrcC-*), were grown at 30°C overnight in Luria Bertani (LB) broth with appropriate antibiotics. The next day, cells were harvested and resuspended in 10 mM MgCl_2_ at OD_600_=0.2 (1 x 10^8^ colony forming units (CFU)/mL). Immediately before inoculations, 0.04% of Silwet L-77 (ThermoFisher Scientific, Waltham, MA) was added to the suspension, and the unifoliate leaves of 14-day-old soybeans were spray-inoculated on adaxial and abaxial surfaces until fully wet. Leaf discs were extracted from 24 plants per treatment, between 1- and 7-days post-infection (dpi). Leaf discs from three plants were pooled per sample, and were used to quantify CFU/cm^2^ as previously described (Liu *et al*., 2015). For phytohormone analysis, eight plants per treatment were sampled at 6- and 24-hours post-infection (hpi) and QuantSeq was performed using 6 hpi samples. g_sw_ in response to *Psg* was measured at 1 hpi using 10 plants per treatment. For QuantSeq analyses, leaves were collected from eight *Psg*-infected and mock-infected plants per CO_2_ treatment and chamber, yielding three biological replicates per treatment for QuantSeq analyses (96 samples, QuantSeq dataset 2). Samples were collected between 09:00 and 10:00 h for all replicates.

### Bacterial growth curves

*Pto*DC3000 cultures were inoculated to a starting concentration of 1 x 10^7^ CFU/mL in 100 mL of LB broth containing appropriate antibiotics. Cultures were grown in temperature-controlled growth chambers maintained at 28°C (no light), ∼20% relative humidity, and either 419 or 550 ppm of CO_2_. Cultures were shaken at 100 rpm and OD_600_ was recorded every four hours over 32 hours.

### QuantSeq

Total RNA was extracted from 50 mg of soybean leaf tissue using the Direct-zol RNA Miniprep Plus Kit with DNaseI treatment (Zymo Research, Irvine, CA). RNA samples were quantified using a Qubit 4.0 Fluorometer (Invitrogen, Carlsbad, CA), and RNA integrity was assessed using the Fragment Analyzer Automated CE System. All RNA samples used for sequencing had an RQN (RNA Quality Number) equal to or greater than 7. Library preparation was performed using the QuantSeq 3’ mRNA-Seq Library Prep Kits (Lexogen, Greenland, NH), and samples were sequenced using the Illumina NovaSeq 6000 System (Iowa State University DNA Facility).

### Bioinformatic and Statistical Analysis of QuantSeq data

For each of the QuantSeq datasets (48 and 96 libraries), libraries were each sequenced on two lanes, generating two sequence files per library. Files were inspected with FASTQC (de Sena Brandine & Smith, 2019) to confirm sequence quality and quantity. Trimmomatic was used to remove adaptor sequences, low quality bases and reads less than 50 bp in length (Bolger *et al*., 2014). STAR version 2.7.10 (Dobin *et al*., 2013) was used to align reads to the Williams 82 reference genome sequence (Wm82.a4, (Schmutz *et al*., 2010)). Samtools version 1.17 (Li *et al*., 2009) was used to merge read data from separate lanes and identify uniquely mapping reads within the combined mapping files (bam files). Mapping files were imported into RStudio version 4.2.3 (Dobin *et al*., 2013) using Rsamtools (Morgan *et al.,* 2021). The gene feature file (GFF) corresponding to the reference genome was imported using rtracklayer (Lawrence *et al*., 2009). The number of reads per sample aligning to each gene was counted using summarizeOverlaps and a count table for all expressed genes within a tissue was generated. Over 329 million and 659 million uniquely mapping reads were identified across QuantSeq datasets 1 and 2, respectively. Data were normalized using the Trimmed Mean of M values (Gupta *et al*., 2021) in the Bioconductor package edgeR (Robinson & Smyth, 2007; Robinson *et al*., 2010; McCarthy *et al*., 2012; Zhou *et al*., 2014). Only genes with log2 counts per million (cpm) > 1 in at least three replicates were used in the analysis. ggplot2 (Wickham, 2009) was used to generate principal component, biological coefficient of variance, and MA plots to visually compare sample replicates and ensure reproducibility (Dziuda, 2010). Two and eight libraries with less than 1 million uniquely mapping reads were removed from subsequent analyses from QuantSeq datasets 1 and 2, respectively, leaving a minimum of six plant replicates per condition. Following renormalization, edgeR was used to analyze differentially expressed genes (DEGs) among the different datasets. For QuantSeq dataset 1, we identified DEGs whose expression changed in response to CO_2_ levels (*e*CO_2_ levels/*a*CO_2_) at 21 days using a false discovery rate (FDR) < 0.01. For QuantSeq dataset 2, we identified DEGs responding to *Psg* treatment (*Psg*/mock) in control and *e*CO_2_ conditions and DEGs whose expression changed in response to CO_2_ in *Psg*-treated (*Psg e*CO_2_/*Psg* control CO_2_) or mock-treated samples (Mock *e*CO_2_/*a*CO_2_) using an FDR < 0.01. Hierarchical clustering in RStudio using hclust (Murtagh, 1985) was used to group similarly expressed genes from DEG lists of interest. Heatmaps were generated using Heatmap.2 in ggplot2 (Wickham, 2009).

### DEG Annotation and Analysis of Overrepresented Transcription Factors and Promoters

DEGs were annotated using the SoyBase Gene Annotation Lookup Tool for Wm82.a4 (https://www.soybase.org/genomeannotation/). For given DEGs of interest, the best BLASTP Arabidopsis hits (TAIR version 10) from the annotation were used to query the STRING database (Szklarczyk *et al*., 2023). To identify transcription factors (TFs) within DEG lists of interest, we used custom perl scripts to query *G. max* transcription factor information downloaded from the Plant Transcriptional Regulatory Map/Plant Transcription Factor Database (PlantRegMap/PlantTFDB v5.0, (Tian *et al*., 2020), https://plantregmap.gao-lab.org).

To identify TF binding sites significantly overrepresented within promoters of DEGs of interest we used the PlantRegMap TF enrichment tool (https://plantregmap.gao-lab.org/tf_enrichment.php) for *G. max*, using all available methods and threshold p-value < 0.05.

### Single Phase Extraction for JA-Targeted and SA (Non-Targeted) Metabolomic Analyses

Approximately 100 mg of each sample was ground and spiked with an internal standard containing 10 µg nonadecanoic acid (1 mg/mL in ethanol) and JA-d5 (Cayman Chemical, Ann Arbor, MI) (JA-d5) (2.5 µg/mL in ethyl acetate). The extraction was initiated with the addition of 90% ice cold methanol using a modified version of the methanolic extraction protocol established previously (Jiye *et al*., 2005).

### GC-MS Acquisition and Analysis for SA

Six hundred microliters of the combined extracts were aliquoted and dried using a speed-vac concentrator for 10 hours and then derivatized according to established protocols (Koek *et al*., 2006). Gas Chromatography Mass Spectrometry (GC-MS) analysis was performed with an Agilent 6890 gas chromatograph coupled to a model 5973 Mass Selective Detector (Agilent Technologies, Santa Clara, CA) using a HP-5MSI 5% phenyl methyl silox with 30 m × 250 µM × 0.25 µm film thickness (Agilent Technologies). Identification and quantification were conducted using AMDIS (Automated Mass spectral Deconvolution and Identification System, National Institute of Standards and Technology, Gaithersburg, MD) with a manually curated, retention indexed GC-MS library with additional identification performed using the NIST20 and Wiley 11 GC-MS spectral library (Agilent Technologies, Santa Clara, CA). Final quantification was calculated by integrating the corresponding peak areas relative to the area of the nonadecanoic acid internal standard. Raw data were normalized to the amount of tissue used.

### LC-MS Acquisition for Targeted JA Analysis

An Oasis Hydrophilic-Lipophilic-Balanced (HLB) column (30 mg sorbent) (Waters Corporation, Milford, MA) was used for the purification of JA as established previously (Kojima *et al*., 2009; Kojima & Sakakibara, 2012). Liquid chromatography (LC) separations were performed with an Agilent Technologies 1290 Infinity II UHPLC instrument equipped with an Agilent Technologies ZORBAX Eclipse C18 (1.8 μm 2.1 mm × 100 mm) analytical column that was coupled to a 6470 triple quadrupole mass spectrometer with an electrospray ionization source (Agilent Technologies, Santa Clara, CA). A volume of 20 µL of each sample was injected into the LC system with the mass spectrometer in negative mode. Multiple reaction monitoring (MRM) was used for detection with transitions of m/z 209→59 (JA) and 214→62 (JA-d5), with fragmentation conducted at 25 eV for both JA and JA-d5.

### Viral infection assays

Lyophilized soybean leaves infected with BPMV or SMV were ground in 10 mL of 50 mM potassium phosphate buffer (pH 7.5) per 1 g of tissue. One unifoliate leaf of 14 dap soybean was dusted with carborundum (Fisher Scientific, Waltham, MA) and rub-inoculated with 50 μL of inoculum or buffer (mock). The second unifoliate leaf was inoculated two days later using the same method. Systemic leaves were sampled from six plants per treatment, at 14 and 21 dpi to assess viral titer. Fresh weight, dry weight, and gene expression were measured using 21 dpi plants. Disease severity was evaluated between 0 and 21 dpi using a single-digit disease rating scale ranging from 0 to 3 (0 = no disease, 1 = mild, 2 = moderate, and 3 = severe). Area under disease progress curve (AUDPC) was calculated based on the equation below (Simko & Piepho, 2012).

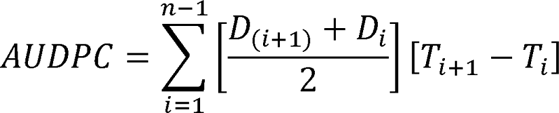

### *F. virguliforme* and *P. sylvaticum* infection assays

Sterilized sorghum (*Sorghum bicolor*) seeds were inoculated with *F. virguliforme* (isolate LL0036) and incubated in the dark at room temperature for 21 days and dried for 3-4 days. Styrofoam cups (8 oz.) were filled with a mixture of 180 mL of sterile sand soil and 9 mL of *F. virguliforme*-infested sorghum. Soybean seeds were placed in the cup and covered with 25 mL of peat mix. At 35 dap, roots and shoots were sampled from six cups of each CO_2_ treatment to measure fresh weight and dry weight. Disease severity was assessed on leaves on a 0 to 7 rating scale as described (Roth *et al*., 2019) (Table S2), and the AUDPC was calculated. For plate growth assays, a starting culture of *F. virguliforme* was prepared on 1/3 strength potato dextrose agar (PDA) and grown at room temperature for 21 days in the dark. A 7-mm plug was punched out of the starting culture and placed into the center of a 90-mm-diameter Petri dish containing 1/3 strength PDA. Ten plates per CO_2_ treatment were maintained in the dark under *a*CO_2_ or *e*CO_2_. Fungal growth was assessed every other day for 11 days.

Millet was inoculated with a 3-day-old culture of *P. sylvaticum* and incubated in the dark at room temperature for 7-14 days (Matthiesen *et al*., 2016). A mixture of 100 mL of sterile sand soil and 5 mL of *P. sylvaticum*-infested millet was placed in a 237 mL (8 oz.) polystyrene cup and covered with 25 mL of peat mix. The seed was placed on top of the peat mix and then covered with another 25 ml layer of peat mix. At 14 dap, plants in six cups of each treatment were up-rooted, roots were washed, and disease severity was assessed using a 0 to 4 rating scale adapted from (Zhang & Yang, 2000) (Table S3). Root and shoot fresh weight and *P. sylvaticum* copy numbers were determined at 21 dap. For plate growth assays, a starting culture of *P. sylvaticum* (isolate Gr8) was grown on 1/2 strength PDA at room temperature for 5 days in the dark. A 7-mm plug was punched and transferred into the edge of a 90-mm-diameter Petri dish containing 1⁄2 strength PDA. Ten plates per treatment were incubated in the dark under *a*CO_2_ or *e*CO_2_. Oomycete growth was measured from the edge of the inoculum plug to the tip of the longest hypha and was assessed every 24 h for 96 h.

### Statistical analysis

Linear mixed-effects models (LMM) were fit to each response following the experimental design. Taking the bacterial infection experiment measuring CFUs as an example, the whole-plot factor was CO_2_ level and the whole-plot experimental units were growth chambers. Randomized complete block design was implemented at the whole-plot level, with each block comprising a pair of chambers at *a*CO_2_ or *e*CO_2_. At the split-plot level, 24 plants were assigned to each pathogen, with sets of three plants pooled for each CFU measurement, repeated over time. Following this split-plot design, the fixed effects of the LMM include the main effects and all interaction effects of the factors CO_2_, bacterial species, and time, and random effects include block, chamber, split-plot, and sets of plants. Similar to this example, LMMs were fit for other responses following the corresponding experimental designs.

For response variables that exhibited unequal variances on the original scale, LMM analysis was done on power- or log-transformed data, and the Delta method was used to compute standard errors for the original scale. For all LMM analysis, model diagnostic checks were done to ensure that model assumptions were appropriate. Parameters of LMM were estimated using lmer() in the lme4 R package. The emmeans() function in the emmeans R package was used to compute the SE. Type III ANOVA tables with F tests were conducted by anova() and the lmerTest R package.

## Results

### Elevated CO_2_ alters soybean physiology and gene expression

Prior to studying the effects of *e*CO_2_ on soybean defense, we wanted to ensure that our growth conditions caused changes in physiological responses that were previously observed under higher atmospheric CO_2_ (Ainsworth & Long, 2021; Li *et al*., 2021). At 14 dap, soybean plants grown in *e*CO_2_ displayed no difference in ΦPSII activity compared to plants grown at *a*CO_2_ (Figure 1a). However, at 21 and 35 dap, ΦPSII activity was higher in *e*CO_2_ (Figure 1a), indicating a higher photosynthetic rate. Stomatal conductance to water vapour (g_sw_) was also affected, with plants grown at *e*CO_2_ displaying lower g_sw_ at 21 and 35 dap (Figure 1b), which was consistent with the lower stomatal density (Figure 1c) and reduced stomatal aperture (Figure 1d) on the abaxial leaf surface. In line with the lower stomatal density, a reduction in stomatal index was observed at *e*CO_2_, indicating that the decrease in stomatal density was due to reduced stomatal production and not an increase in leaf expansion (Figure S2). At 35 dap, soybeans grown in *e*CO_2_ had greater shoot fresh weight (Figure 1e) and dry weight (Figure 1f) and were visibly larger than those grown in *a*CO_2_ (Figure 1g). Together, these experiments demonstrate that a 31% increase in [CO_2_] induced physiological responses under our conditions that were consistent with prior studies.

**Figure 1.**
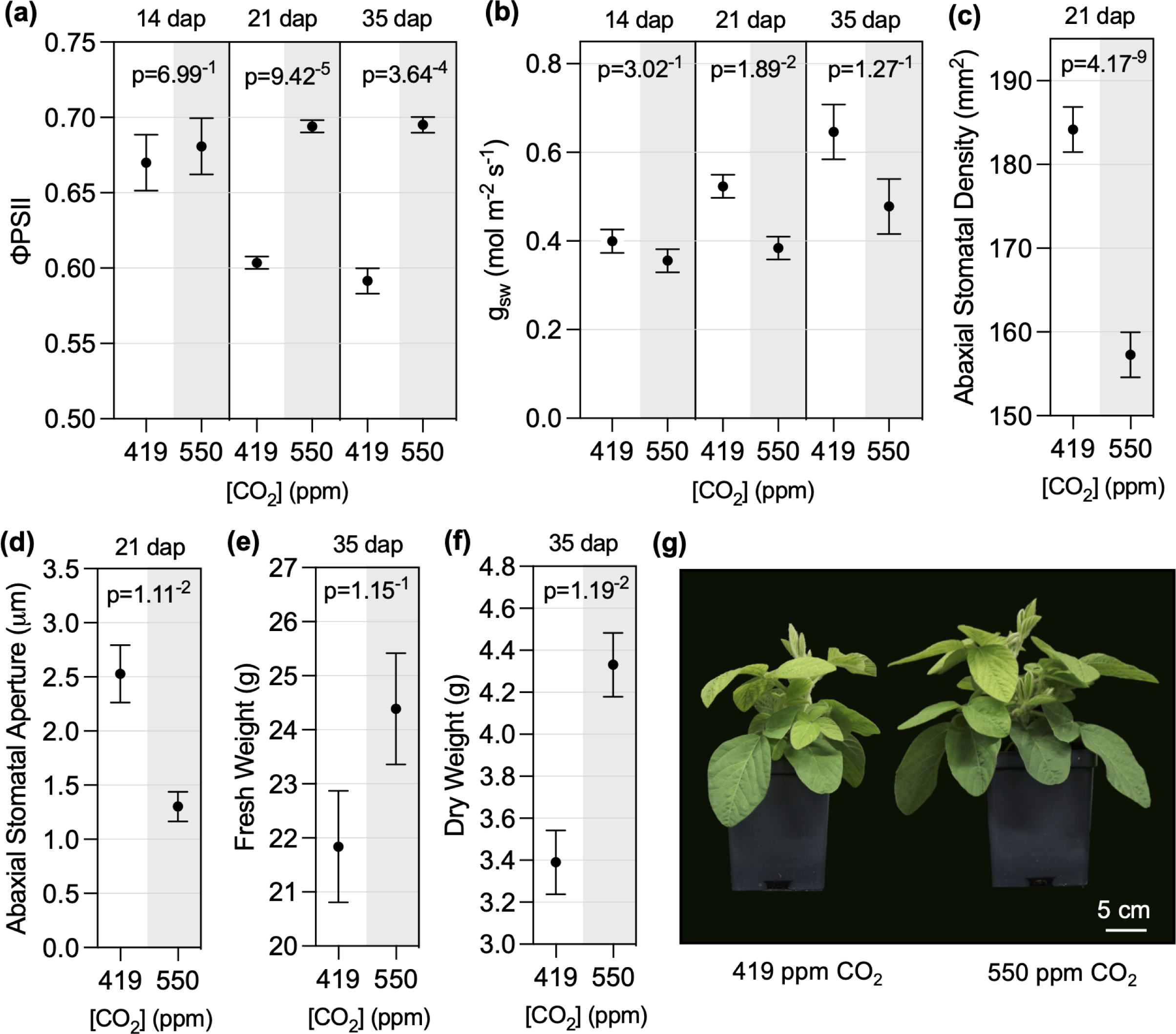
Effects of *e*CO_2_ on soybean physiology and growth. **(a)** Photosystem II (ΦPSII) activity and **(b)** stomatal conductance (g_sw_) were measured at the indicated days after planting (dap) using a LI-600 portable system. **(c)** Stomatal density and **(d)** stomatal aperture measured at 21 dap using three randomly selected fields of view per leaf sample using a brightfield microscope. **(e)** Shoot fresh weight, **(f)** shoot dry weight, and **(g)** representative photos of soybean plants at 35 dap. All measurements and samples were taken from the newest fully expanded leaves of 10 plants at each time point, except ΦPSII and g_sw_, which were measured in 15 plants per CO_2_ treatment. The three experimental replicates were conducted simultaneously in independent CO_2_ controlled chambers using a replicated completely randomized design. Data was graphed as the mean across the three replicates with standard error bars. Linear mixed effect model (LMM) analysis was applied to the fourth power of ΦPSII 35 dap and log-transformed stomatal aperture data due to unequal variance on the original scale. P-values were computed based on F-tests for the effect of CO_2_ at each time point from the LMM analysis.

To investigate gene expression changes associated with physiological responses to *e*CO_2_, we conducted QuantSeq analysis using RNA isolated from unifoliate leaves at 21 dap. We identified 388 DEGs (Table S4), which were used to construct a heat map that formed two distinct clusters associated with [CO_2_] (Figure 2a). The 180 DEGs in cluster 1 were repressed in *e*CO_2_, while the 208 DEGs in cluster 2 were induced in *e*CO_2_ (Table S4). To identify gene networks responding to [CO_2_], the 321 best BLASTP Arabidopsis homologs corresponding to our 388 soybean DEGs were were used as input for STRING (Szklarczyk *et al*., 2023) and gene ontology (GO) analyses (Szklarczyk *et al*., 2023) to identify biological processes (BP) that were overrepresented within each cluster (Table S5). In cluster 1, two major themes emerge related to i) responses to JA signaling (*e.g.* response to wounding, response to hormone, regulation of defense response, response to lipid, regulation of JA-mediated signaling) and ii) photosynthesis (*e.g.* response to light intensity, response to high light intensity, photosynthesis, photosynthesis light reaction). The STRING association network produced from the genes in cluster 1 also highlights the effects on JA signaling with a node of highly connected genes that include *MYC2* transcription factor and *TIFY* (JAZ repressor) genes (Figure 2b, light blue). A node of highly connected genes in the lower right quadrant includes several genes related to photosynthetic processes, including soybean homologs of *AtRbcS1B, AtRAF1, AtLHCB8, AtLHCB1, AtCHLM*, and *AtPIF1* (Figure 2b, light orange).

**Figure 2.**
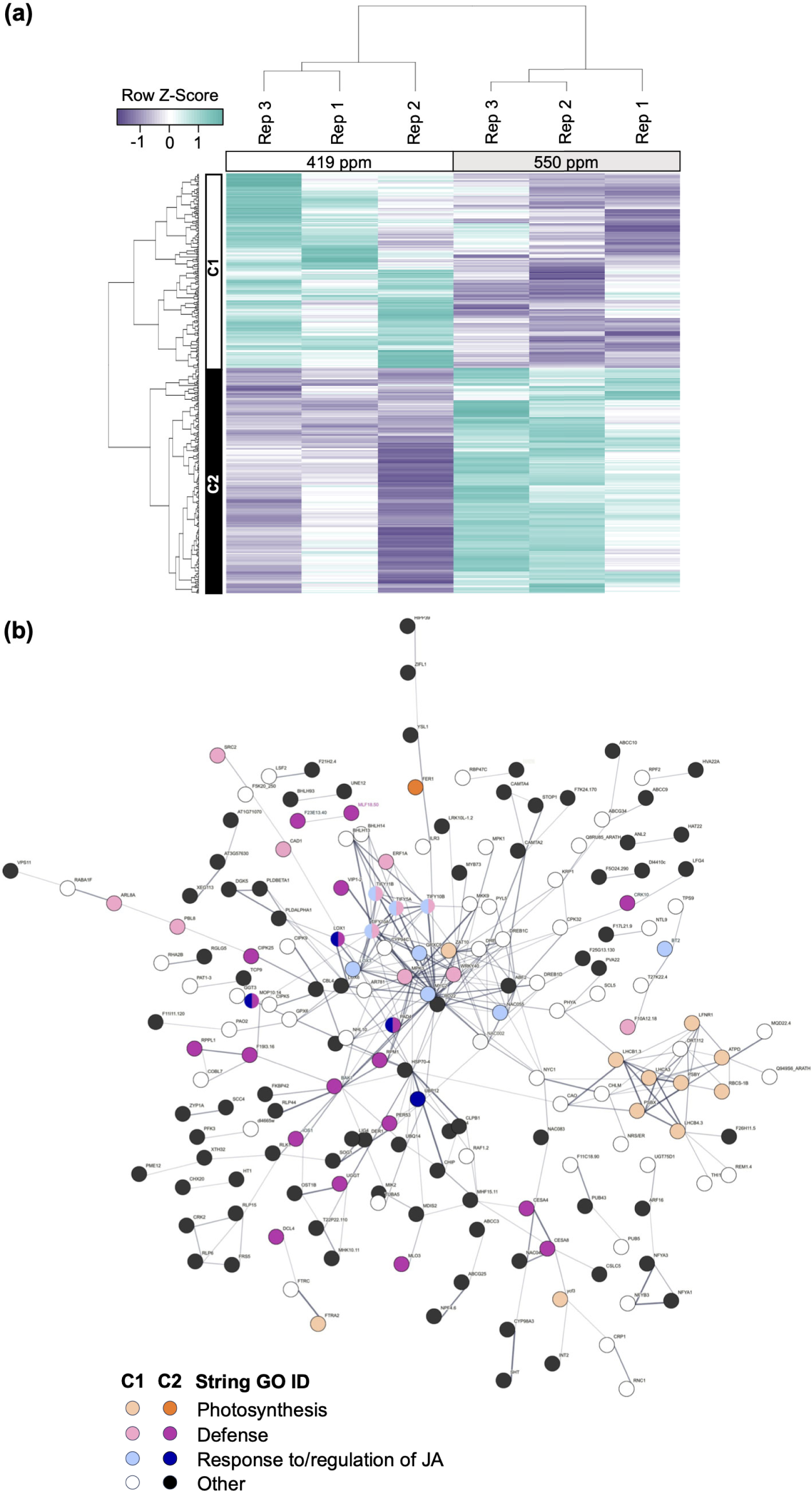
Transcriptomic analysis of soybean gene expression in leaves of plants grown under *a*CO_2_ (419 ppm) or *e*CO_2_ (550 ppm). **(a)** DEGs responding to *e*CO_2_ were identified using a FDR < 0.01 (Table S3). Samples for QuantSeq analysis were taken from the newest fully expanded leaves of soybean plants at 21 days after planting. The three experimental replicates were conducted simultaneously in independent CO_2_ controlled chambers using eight plants per CO_2_ treatment. Row Z-scores were used for hierarchical clustering of DEGs, based on expression across samples and replicates. Two expression clusters were identified. DEGs in cluster 1 were expressed at higher levels in 419 ppm vs 550 ppm CO_2_ and DEGs in cluster 2 were expressed at higher levels in 550 ppm vs 419 ppm CO_2_. **(b)** STRING network for DEGs identified between *a*CO_2_ and *e*CO_2_ in leaves at 21 (dap). The 388 genes identified in soybean correspond to 321 genes in *Arabidopsis thaliana* that were used to assign functional annotations. Lightly shaded or white circles indicate DEGs from cluster 1 (C1) and brightly shaded and black circles indicate DEGs from cluster 2 (C2).

Major themes emerging from cluster 2 are BP terms related to i) defense (*e.g.* defense, response to external stimulus, response to biotic stimulus, response to bacterium, response to other organism) and ii) protein modification (*e.g.* protein phosphorylation, protein modification process, phosphorus metabolic process) (Table S5). The STRING association network identifies interconnected genes associated with defense responses (Figure 2b). These nodes include soybean homologs of *AtBAK1*, *AtPAD4*, and *AtIOS1* (Table S4), all required for pattern-triggered immunity (PTI) responses (Yeh *et al*., 2016; Pruitt *et al*., 2021), and a number of resistance-like proteins, including *AtRPM1*, *AtRLP1*, and *AtRLP44*. While not highlighted in the STRING analyses, soybean homologs of genes associated with reduced g_sw_ and stomatal aperture (*AtHT1*, *AtFER*, *AtZIFL1*, *AtPLDa1*, and *AtNRT1:2*) were upregulated in *e*CO_2_ (Table S4). Together, our gene expression analyses indicate that *e*CO_2_ causes changes in the transcriptome of unifoliate soybean leaves, repressing expression of photosynthetic genes, altering JA signaling, priming innate immunity against microbial pathogens and promoting expression of genes associated with stomatal closure.

### Soybean resistance to *Pseudomonas spp.* is enhanced under *e*CO_2_

In order to assess the impacts of *e*CO_2_ on susceptibility to microbial pathogens, we first used the well characterized soybean-*P. syringae* pathosystem (Lindeberg *et al*., 2009; Whitham *et al*., 2016). Basal defense against bacterial pathogens is initiated by cell surface receptors that recognize microbe-associated molecular patterns (MAMPs), such as the bacterial flagellin peptide flg22 (DeFalco & Zipfel, 2021). *Psg*, which causes bacterial blight in soybean, was chosen to model a compatible interaction where effectors effectively suppress immunity resulting in disease (Prom & Venette, 1997). At 1 dpi, the proliferation of *Psg* was 1.1-log_10_ fold lower in plants grown in *e*CO_2_ compared to *a*CO_2_ (Figure 3a), and it remained significantly reduced in *e*CO_2_ over the 7-d time course (Figure 3a). In compatible interactions, *Psg* infection causes the development of necrotic spots that become surrounded by yellow halos (Budde & Ullrich, 2000). At 7 dpi, soybeans infected under [*a*CO_2_] displayed necrotic lesions across the surface of leaves, consistent with bacterial blight symptoms (Figure 3b). In contrast and consistent with reduced *Psg* growth, few disease symptoms were observed on the leaves of plants grown at *e*CO_2_ (Figure 3b).

**Figure 3.**
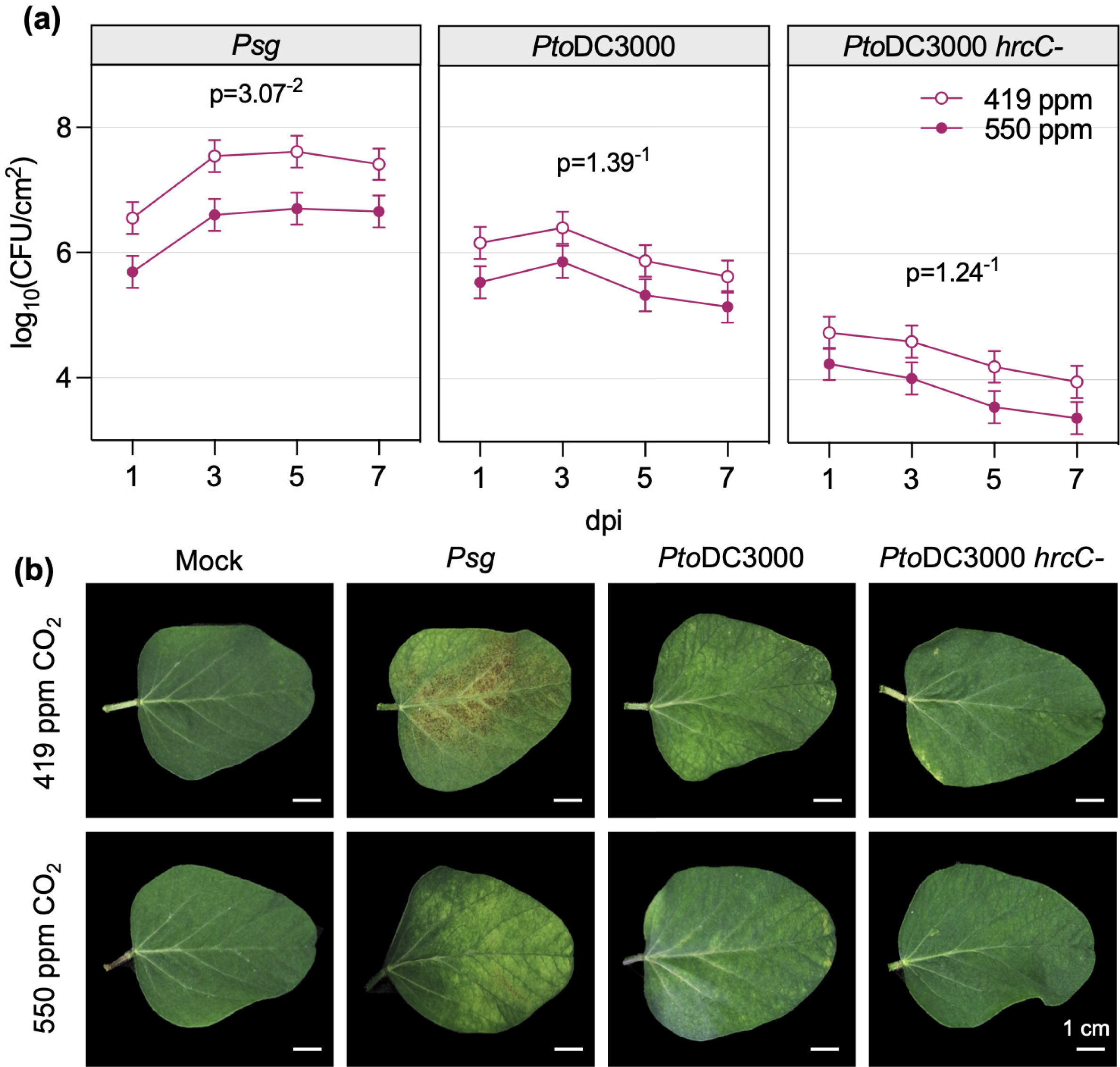
Soybean plants growing in *e*CO_2_ are less susceptible to *P. syringae*. **(a)** Colony forming units (CFU) of *P. syringae* pv. *gycinea* (*Psg*), *P. syringae* pv. *tomato* (*Pto*DC3000) and *Pto*DC3000 *hrcC^-^*were quantified in the unifoliate leaves of plants at 1, 3, 5, and 7 days post-inoculation (dpi). The unifoliate leaves of 24 plants were sampled for each treatment group, and three independent biological replicates were performed. Data points represent the mean CFU count of the three biological replicates with standard error bars. P-values were computed using t-tests for the contrast between two CO_2_ levels of each pathogen averaged over dpi from the LMM analysis. **(b)** Representative images of unifoliate leaves photographed at 7 dpi. Mock plants were treated with 10 mM MgCl_2_, 0.04% Silwet L-77 solution with no bacteria.

We also inoculated plants with *Pto*DC3000, as an example of an incompatible interaction causing hypersensitive cell death resulting in disease resistance (Whalen *et al*., 1991), and the *Pto*DC3000 type III secretion mutant, *hrcC-*, that cannot secrete bacterial effector proteins and therefore cannot multiply or induce hypersensitive cell death (Brooks *et al*., 2004). Lower CFUs were consistently observed across the time course in soybean grown at *e*CO_2_ when inoculated with either *Pto*DC3000 or *hrcC-*, although the p-values were greater than 0.05 (Figure 3a). At 7 dpi, *Pto*DC3000-infected plants grown at either [CO_2_] began to show signs of a hypersensitive reaction while *hrcC-* infection did not result in obvious disease or defense phenotypes, regardless of the [CO_2_] (Figure 3b).

To assess if atmospheric [CO_2_] independently affects bacterial growth rate, we conducted a growth curve analysis on liquid cultures of *Pto*DC3000 in our controlled environment growth chambers. The growth curves of *PtoDC3000* cultures were similar between the two CO_2_ levels (Figure S3), indicating that reduced growth of *P. syringae* in soybean is not due to direct effects of *e*CO_2_ on bacterial multiplication. Together, these data suggest that *e*CO_2_ enhances resistance to *P. syringae* infection in soybean during compatible interactions and, to a lesser extent, during an incompatible interaction.

### Elevated CO_2_ alters basal immune responses

To determine if *e*CO_2_ affects immunity to *Psg* on a molecular level, we assessed multiple hallmarks of plant immune signaling. In response to *Psg*, the change in stomatal conductance following *Psg* treatment did not differ between [CO_2_] (Figure 4a). However, soybeans grown at *e*CO_2_ exhibited greater overall production of reactive oxygen species when considering both mock- and flg22-treated leaf discs (Figure 4b) as well as higher and more sustained MAPK activation (Figure 4c). We also observed that a soybean *PR1* homolog, an SA-regulated defense gene, accumulated to higher levels in mock- and *Psg*-inoculated plants grown in *e*CO_2_, particularly in the 24 hpi sample (Figure 4d,e) and corresponded to a slightly greater SA accumulation in leaves sampled at 6 hpi (Figure S4a). At 24 hpi, expression of the JA marker gene *KTI1* was more down regulated in response to *Psg* (Figure 4f,g). However, JA accumulation was not affected by [CO_2_] in our metabolomics analysis (Figure S4b). Together, these data suggest that bacterially-induced immune signaling is more robust in plants grown in *e*CO_2_, which correlates with the enhanced resistance to *Pseudomonas spp.* in these plants.

**Figure 4.**
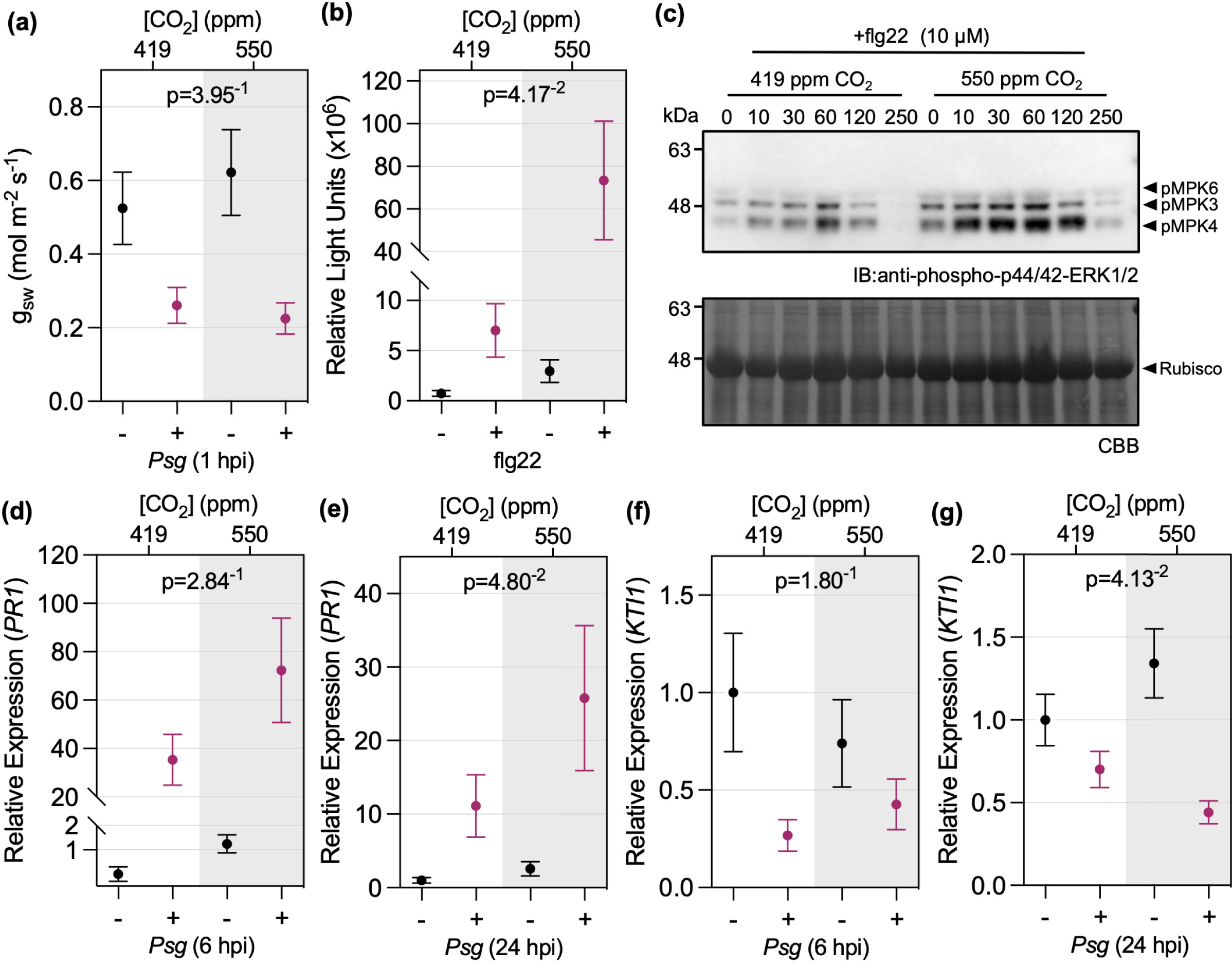
Bacterially-induced defense signaling is more strongly upregulated in soybean leaves grown under *e*CO2. **(a)** Stomatal conductance (g*sw*) at 1 hour post-inoculation (hpi) with *P. syringae* pv. *glycinea* (*Psg*). **(b)** Reactive oxygen species (ROS) production in response to bacterial flagellin 22 (flg22) peptide was quantified using a chemiluminescence assay. Relative light units were determined using leaf discs from 12 plants per treatment, and **(c)** flg22-induced MAPK activation using protein extracted from 12 plants per time point (0 to 250 minutes after treatment) and visualized by immunoblot analysis. Expression of *PR1* (SA marker gene) **(d)** 6 hpi and **(e)** 24 hpi and expression of *KTI1* (JA marker gene) **(f)** 6 hpi and **(g)** 24 hpi with *Psg* or mock treatment was assessed by RT-qPCR. RNA was extracted from six plants per treatment and expression was measured relative to the *Skp1* housekeeping gene. All experiments were conducted using leaves or leaf discs collected from 14-day old plants grown under 419 ppm or 550 ppm CO_2_. Three independent replicates were conducted for each experiment. Data points represent average values across the three experimental replicates with standard error bars. P-values for ROS production and *PR1* expression were computed using F-tests for the main effect of CO_2_, and g*sw* and *KTI1* expression using the interaction effect between CO_2_ and *Psg*, from the LMM analysis based on log-transformed data.

### *e*CO_2_ alters the soybean transcriptome in response to *Psg*

To gain further insight into the effects of *e*CO_2_ on molecular responses to *Psg* treatment, we conducted QuantSeq analysis on soybean leaves at 6 hpi with buffer alone (mock) or with *Psg*. We identified DEGs in mock-versus *Psg-*treated plants grown at each of the two [CO_2_]: 18,054 DEGs were identified in plants grown in *a*CO_2_ and 19,443 DEGs were identified for plants grown in *e*CO_2_. We produced a combined list of all unique DEGs for a total of 21,822 *Psg-* responsive DEGs, which was used to generate a heat map that formed four distinct clusters (Figure S5; Table S6). Cluster 1 (1,661 DEGs) had mixed expression in response to *Psg* treatment and no significantly (corrected p<0.05) overrepresented GO terms (Table S7). Cluster 2 (9,822 DEGs) was induced by *Psg* treatment and had 81 significant GO terms associated with stress (response to cadmium ion, heat, hypoxia, oxidative stress, salt, and UV-B), defense (response to chitin, defense response to bacterium, fungus, and virus, cell death, regulation of SA biosynthesis, response to SA, hypersensitive response, and systemic acquired resistance), and ER stress (protein folding, response to unfolded protein, and response to ER stress). Clusters 3 (6,718 DEGs) and 4 (3,621 DEGs) were repressed by *Psg* treatment. While cluster 3 had 50 significant GO terms associated with photosynthesis, response to light, and circadian rhythm, cluster 4 had two significant GO terms: response to starvation and oxidation-reduction process. Among DEGs repressed by *Psg* treatment in *a*CO_2_ and *e*CO_2_ conditions, 61.6% had greater negative fold-changes in *e*CO_2_ (7,419/12,053 DEGs). Similarly, for DEGs induced by *Psg* in *a*CO_2_ and *e*CO_2_ conditions, 60.1% had greater positive fold changes in soybeans grown in *e*CO_2_ (5,671/9,357 DEGs).

To further understand how responses to *Psg* differ with changes in atmospheric [CO_2_], we identified 622 and 1,919 DEGs associated with *e*CO_2_ versus *a*CO_2_ in mock- or *Psg*-inoculated plants, respectively (Table S8). Using a union of these two lists, we generated a heatmap of the 2,419 DEGs, which formed five distinct clusters (Figure 5). To assign potential function to each of the clusters, we used the Arabidopsis best BLASTP homologs corresponding to all DEGs within a cluster as input into STRING (Szklarczyk *et al*., 2023) and performed GO BP enrichment for each cluster (Table S9). Cluster 1, which tended to be induced by *e*CO_2_ in mock and *Psg* samples, was associated with 40 GO terms related to stress (response to chemical, stress, and abiotic stimulus) and defense (regulation of JA signaling, response to JA, response to bacterium, and defense response). DEGs in cluster 2 were induced by *Psg* with greater induction observed in *e*CO_2_ and corresponded to 139 significant GO terms related to stress and defense responses. DEGs in cluster 3 were repressed by *e*CO_2_ and corresponded to 37 significant GO terms, many associated with stress and regulation (regulation of transcription, circadian rhythm, primary metabolism, development, and photoperiodism). DEGs in cluster 4 were induced by *Psg* with greater induction observed in *a*CO_2_ and corresponded with 83 significant GO terms associated with stress, signaling, development, and defense. DEGs in cluster 5 were repressed by *Psg*, with greater repression at *e*CO2. The 37 GO terms in cluster 5 were largely associated with photosynthesis and response to light.

**Figure 5.**
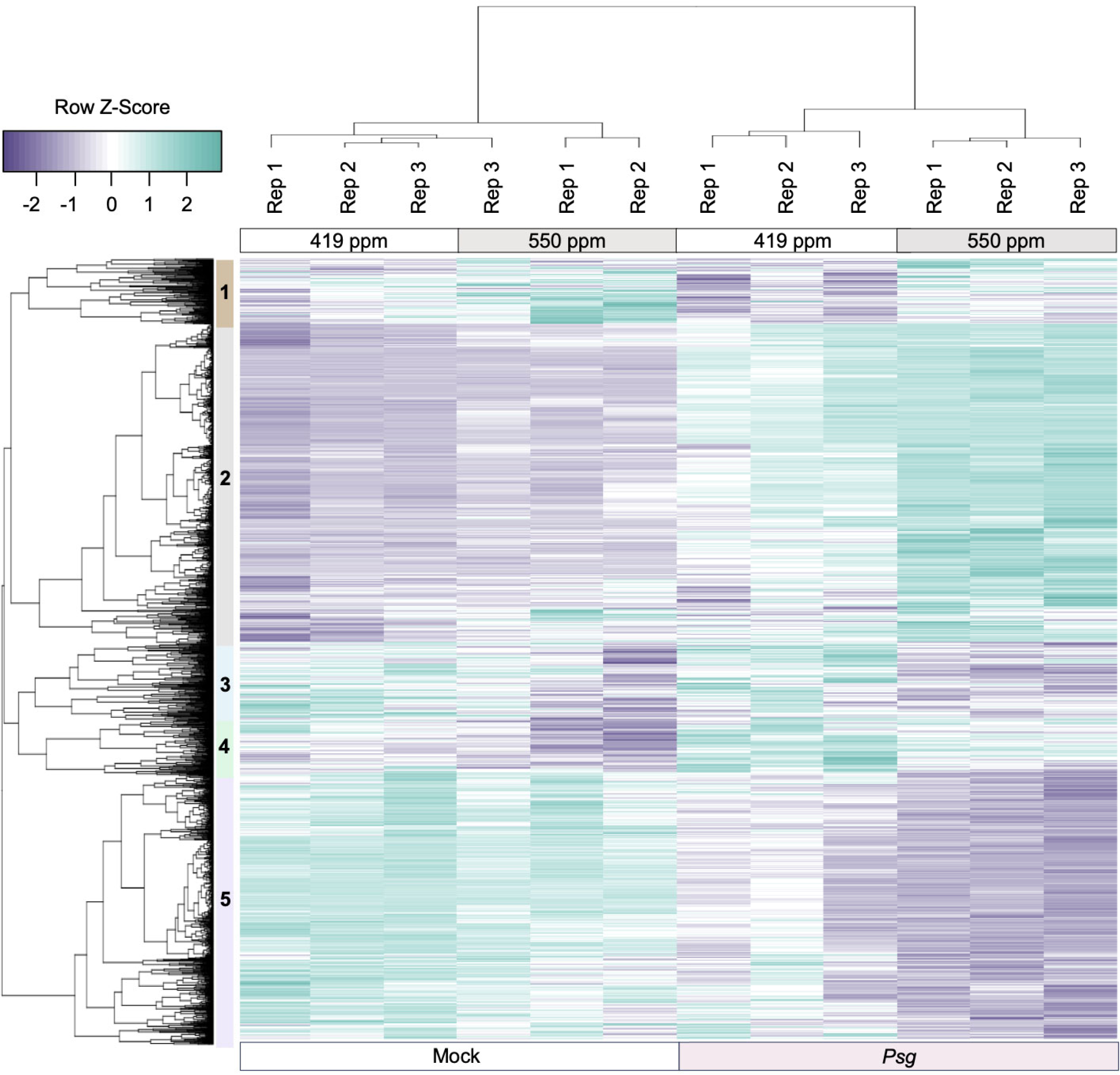
Identification of soybean DEGs responding to eCO_2_ (419 ppm versus 550 ppm) in either mock or *Pseudomonas syringae* pv. *gylcinea (Psg*) infected samples at 6 hpi. Samples for QuantSeq analysis were taken from the unifoliate leaves of 14-day old plants at 6 h post-inoculation with mock treatment or *Psg*. The three independent replicates were conducted using eight plants per CO_2_ treatment. **(a)** Using an FDR < 0.01, we identified 2,419 DEGs responding to eCO2 in *Psg* and/or mock-treated samples. Row Z-scores were used for hierarchical clustering of DEGs, based on expression across samples and replicates. Purple indicates expression values below the row mean for a given DEG and sample and teal indicates expression values above the row mean for a given DEG and sample. Rep indicates the independent biological replicate. The five major expression clusters of DEGs are indicated as 1 to 5, with clusters 2 and 5 containing the most genes. Cluster 2 contains 986 genes that are primarily upregulated at 6 hpi with *Psg* but these genes are upregulated more in *e*CO_2_. Cluster 5 contains 828 genes that are downregulated at 6 hpi with *Psg*, but these genes are more down regulated in *e*CO_2_.

Across all 2,419 DEGs, 74.8% and 83.2% of DEGs responding to *e*CO_2_ in *Psg* or mock were also significantly differentially expressed in response to *Psg* at *a*CO_2_ and eCO_2_, respectively. This confirms that *e*CO_2_ impacts the expression of genes involved in pathogen defense responses. This is most obvious in clusters 2 (986 DEGs) and 5 (828 DEGs), where 71.1% and 94.3% of DEGs are also significantly differentially expressed in response to *Psg* at *a*CO_2_ and 82.3% and 99.5% of DEGs are also significantly *Psg* responsive in *e*CO_2_. In summary, the QuantSeq analyses indicate that *e*CO_2_ does not cause a reprogramming of responses to *Psg*, but rather it enhances the up- or down-regulation of many genes associated with defense and stress responses and photosynthetic processes.

### Identification of regulatory elements controlling response to *e*CO_2_

We next sought to identify the TFs that were differentially expressed in the 2,419 DEGs responding to *e*CO_2_ in Figure 5. Across all five clusters, we identified 144 TFs representing 31 transcription factor families. Clusters 1-5 contained 7, 43, 27, 19, and 48 unique TFs, respectively (Table S10). Given that many of the GO term descriptions significantly overrepresented in clusters 1 through 5 included the words “response to” or “regulation of”, we were interested in predicting upstream TFs that could also be important in *e*CO_2_ responses. Using the DEG list for each cluster as input into PlantRegMap (Tian *et al*., 2020), we identified 1, 44, 18, 13, and 16 significantly overrepresented TF binding sites (TFBS) for clusters 1-5, respectively (Table S11). Interestingly, some predicted TFBS were significant in multiple clusters. In addition, five predicted TFBS were also among our DEGs of interest. We identified the DEG targets corresponding to each overrepresented TFBS (Table S12). In order to assign a possible function to a TF with an overrepresented TFBS, we used the SoyBase GO Term Enrichment Tool (https://www.soybase.org/goslimgraphic_v2/dashboard.php) to identify significantly overrepresented GO terms (corrected p<0.05) associated with the corresponding target DEGs (Table S13). Of the 92 significant TFBS predicted, 48 had significantly overrepresented GO terms. Identified GO terms were associated with stress (response to heat, H_2_O_2_, redox state, phosphate starvation, and light), regulation (cell aging, circadian rhythm, growth rate, proton transport, and transcription), photosynthesis (light reaction, electron transport, and plastid organization), and defense (fatty acid elongation, biosynthesis of anthocyanins, carotenoids, and flavonoids, and negative regulation of defense response to bacterium). Sorting the data in Table S13 revealed multiple TFs were regulating DEGs corresponding to the same GO term. For example, we identified nine significant TFBS associated with photosynthetic electron transport in photosystem I.

Given these results, we wanted to examine the TF/TFBS data relative to our DEGs from mock and *Psg*-infected tissues. We plotted the number of target DEGs corresponding to each predicted TFBS (Figure 6a). For almost every predicted TFBS, there were more targets among DEGs from *Psg*-infected tissues than mock-infected tissues, suggesting *Psg* infection resulted in a more robust response to *e*CO_2_. To confirm this pattern was not due to errors in predicting TFBS, we also plotted the number of DEGs associated with significantly overrepresented GO terms (Figure 6b; Table S8,S9). For almost every significant GO term, there were more GO terms associated with *Psg*-infected tissues than mock-infected tissues. To better visualize the interaction between TF and TFBS, we used the best Arabidopsis homolog of each TF and TFBS as input into STRING (Szklarczyk *et al*., 2023). In Figure 6c, TF and TFBS associated with *Psg*-infected tissues are colored shades of teal, while TF and TFBS associated with mock-infected tissues are colored shades of purple. As in 6a and 6b, the TF and TFBS from *Psg*-infected tissues are far more robust, confirming *Psg*-infection enhances responses to *e*CO_2_.

**Figure 6.**
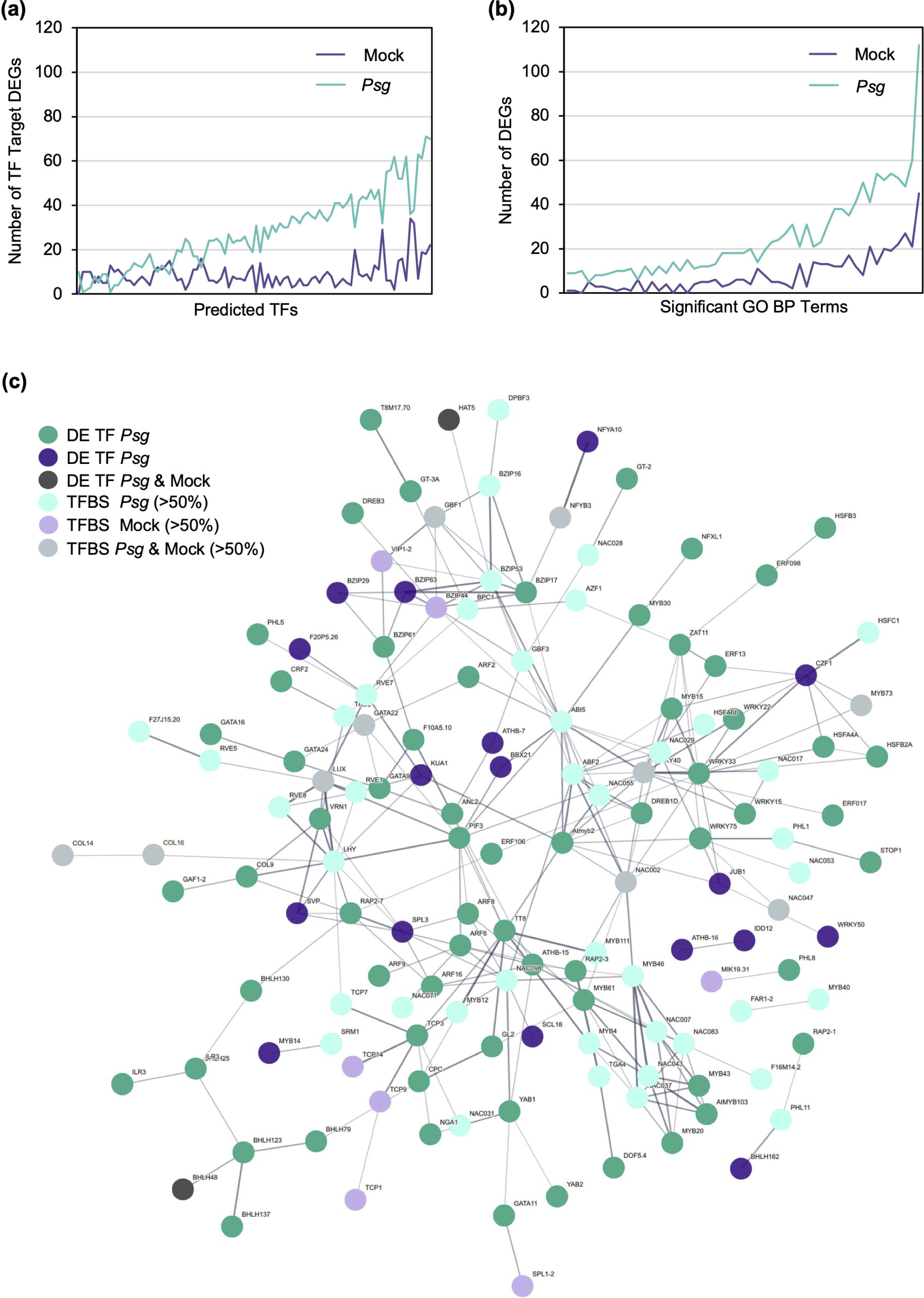
Significantly differentially expressed TFs and over-represented TFBS are more robust in *Psg-*infected tissues responding to eCO_2_ than mock infected tissues. PlantRegMap/PlantTFDB v5.0 <u>(Tian *et al*., 2020)</u> was used to identify significantly differentially expressed TFs and significantly overrepresented TFBS among the DEG clusters identified in Figure 5 (Supplemental Tables S9-S11). **(a)** Significant TFBS versus predicted number of DEG targets. Overrepresented TFBS and DEGs targets are plotted for every cluster (cluster information not shown), therefore if a TFBS was significant in multiple clusters, it was plotted multiple times but with different DEG targets. **(b)** Significantly overrepresented GO terms (minimum of 10 DEGs per GO term) versus DEG number (Supplemental Table S5). In both panels, DEGs in teal were significant in *Psg* samples and/or DEGs in purple were significant in mock treated samples. **(c)** Best Arabidopsis homologs of TFs and TFBS were input into STRING <u>(Szklarczyk *et al*., 2023)</u>. TFs differentially expressed in response to *e*CO_2_ in *Psg*-treated samples, mock-treated samples or both are colored dark teal, dark purple or grey, respectively. TFBS are colored based on the origin of the target DEGs.

### *e*CO_2_ suppresses viral immunity

To determine if *e*CO_2_ affects virus susceptibility in soybean, we inoculated plants with two virus species, BPMV, a *Comovirus* in the family *Secoviridae* (Sanfaçon *et al*., 2009), and SMV, a *Potyvirus* in the family *Potyviridae* (Hajimorad *et al*., 2018). BPMV and SMV caused a greater reduction in soybean growth and shoot dry weight when compared to the mock plants in *e*CO_2_ than in *a*CO_2_ (Figure 7a, S6). Accumulation of BPMV and SMV was higher in the newest fully expanded leaves of infected plants grown at *e*CO_2_ compared to those grown at *a*CO_2_ at 14 dpi (Figure 7b,c) and to a lesser extent at 21 dpi (Figure S7). Accordingly, the AUDPC was greater for BPMV- and SMV-infected plants grown at *e*CO_2_ (Figure 7d,e). We used RT-qPCR to assay expression of soybean homologs of four genes associated with defense responses to viruses: *PR1*, *KTI1*, *AGO1*, and *DCL2*. We found that the overall mean expression of the SA marker gene, *PR1*, was lower at *e*CO_2_ when considering both mock and virus-infected plants (Figure 7f). In addition, expression of *PR1* was induced in both BPMV- and SMV-infected plants relative to mock-inoculation under *a*CO_2_. However, *PR1* expression in the virus-infected plants was similar to the mock-inoculated plants in *e*CO_2_. The expression of *KTI1* was similar under the two CO_2_ levels (Figure 7g), and its induction in virus-infected versus mock plants was not significantly different between *a*CO_2_ and *e*CO_2_, suggesting that JA-mediated defenses against viruses were not affected. We also observed that *AGO1* and *DCL2* accumulated to lower levels in plants grown at *e*CO_2_ (Figure 7h,i), suggesting that RNA silencing-based defenses could potentially be compromised (Leonetti *et al*., 2021; Akbar *et al*., 2022). These data suggest that the increased susceptibility of soybean to BPMV and SMV under *e*CO_2_ may be partially mediated by suppression of SA-related signaling and RNA silencing mechanisms.

**Figure 7.**
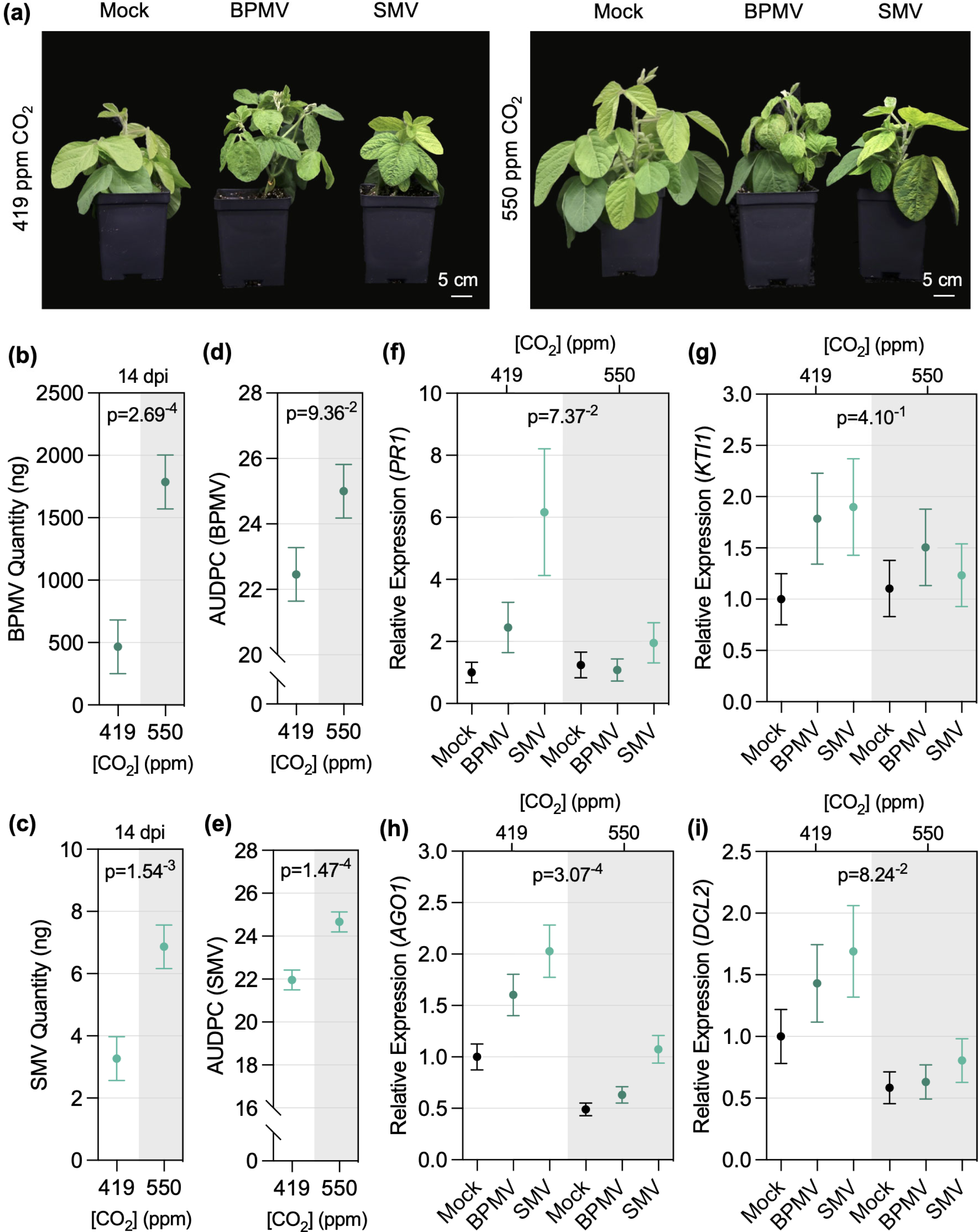
Young soybean plants growing in *e*CO_2_ are more susceptible to BPMV and SMV. **(a)** Representative photos of soybean growth in response to *e*CO_2_ during infection with BPMV or SMV compared to mock treated control plants. Plants were photographed at 21 days post-inoculation (dpi) **(b)** BPMV and **(c)** SMV quantity at 14 dpi was assessed by RT-qPCR using *Skp1* as a housekeeping gene. Disease progression of **(d)** BPMV and **(e)** SMV calculated as the area under the disease progress curve (AUDPC) over a 35 day time course. Expression of **(f)** *PR1* (SA marker gene) **(g)** *KTI1* (JA marker gene), and RNA silencing-related genes **(h)** *AGO1* and **(i)** *DCL2* in response to BPMV, SMV, or mock treatment was assessed by RT-qPCR analysis relative to the expression of *Skp1*. The newest fully expanded leaves of eight plants were sampled at the indicated time point for each treatment. The three experimental replicates were conducted simultaneously in independent CO_2_ controlled chambers using a replicated complete randomized design. Data was graphed as the mean across the three replicates with standard error bars. P-values were computed using F-tests for the main effect of CO_2_ from the LMM analysis on the log-transformed relative gene expression data. The following p-values were associated with the log-transformed relative *PR1* expression of virus-infected plants compared to mock-treated plants at *a*CO_2_ condition: p(BPMV_PR1,419_)=0.1282, p(SMV_PR1,419_)=0.0178 and at *e*CO_2_ condition: p(BPMV_PR1,550_)=0.7779, p(SMV_PR1,550_)=0.3882. All were calculated based on t-tests for the contrasts between each pathogen and mock at the corresponding CO_2_ conditions from the LMM analysis. The following p-values were computed using t-tests from the LMM analysis for the interaction effect between CO_2_ condition and pathogen treatment (virus or mock) on log-transformed *KTI1* gene expression: p(BPMV_KTI1_)=0.6156 and p(SMV_KTI1_)=0.3444.

### Susceptibility to soil-borne filamentous pathogens is altered under elevated CO_2_

To determine if *e*CO_2_ also impacts susceptibility to soil-borne pathogens, we inoculated soybean with *F. virguliforme*, a hemibiotrophic fungus causing sudden death syndrome (SDS) (O’Donnell *et al*., 2010), or *P. sylvaticum*, a necrotrophic oomycete causing seed decay and seedling root rot (Rojas *et al*., 2017). Root and shoot dry weight were decreased to a greater extent in *F. virguliforme-*infected plants grown in *e*CO_2_ compared to plants grown under *a*CO_2_ at 35 dap (Figure 8a,b, S8). Correspondingly, progression of the foliar symptoms of SDS was greater (Figure 8c) and more severe symptoms developed on the leaves under *e*CO_2_ (Figure 8d). Consistent with the increased disease symptoms, *F. virguliforme* was more abundant in the tap roots of plants grown in *e*CO_2_ at 35 dap (Figure 8e). To investigate if expression of SA- or JA- related defense genes was affected in *e*CO_2_, accumulation of *PR1* and *KTI1* mRNA transcripts, respectively, were quantified in soybean roots using RT-qPCR. Expression of neither gene was convincingly altered in response to *F. virguliforme* infection in *e*CO_2_ relative to plants grown under *a*CO_2_ (Figure 8f, S9). To investigate if *F. virguliforme* growth is affected by [CO_2_], we measured the diameter of *F. virguliforme* colonies as they grew on plates for 11 days in *a*CO_2_ or *e*CO_2_. Fungal growth was similar at the two CO_2_ levels (Figure S10), indicating that the greater fungal accumulation in roots and SDS progression observed in soybeans at *e*CO_2_ is not caused by increased pathogen replication rate. Together, our data suggest that *e*CO_2_ increases susceptibility to *F. virguliforme*, although the underlying molecular mechanisms mediating this altered interaction are unclear.

**Figure 8.**
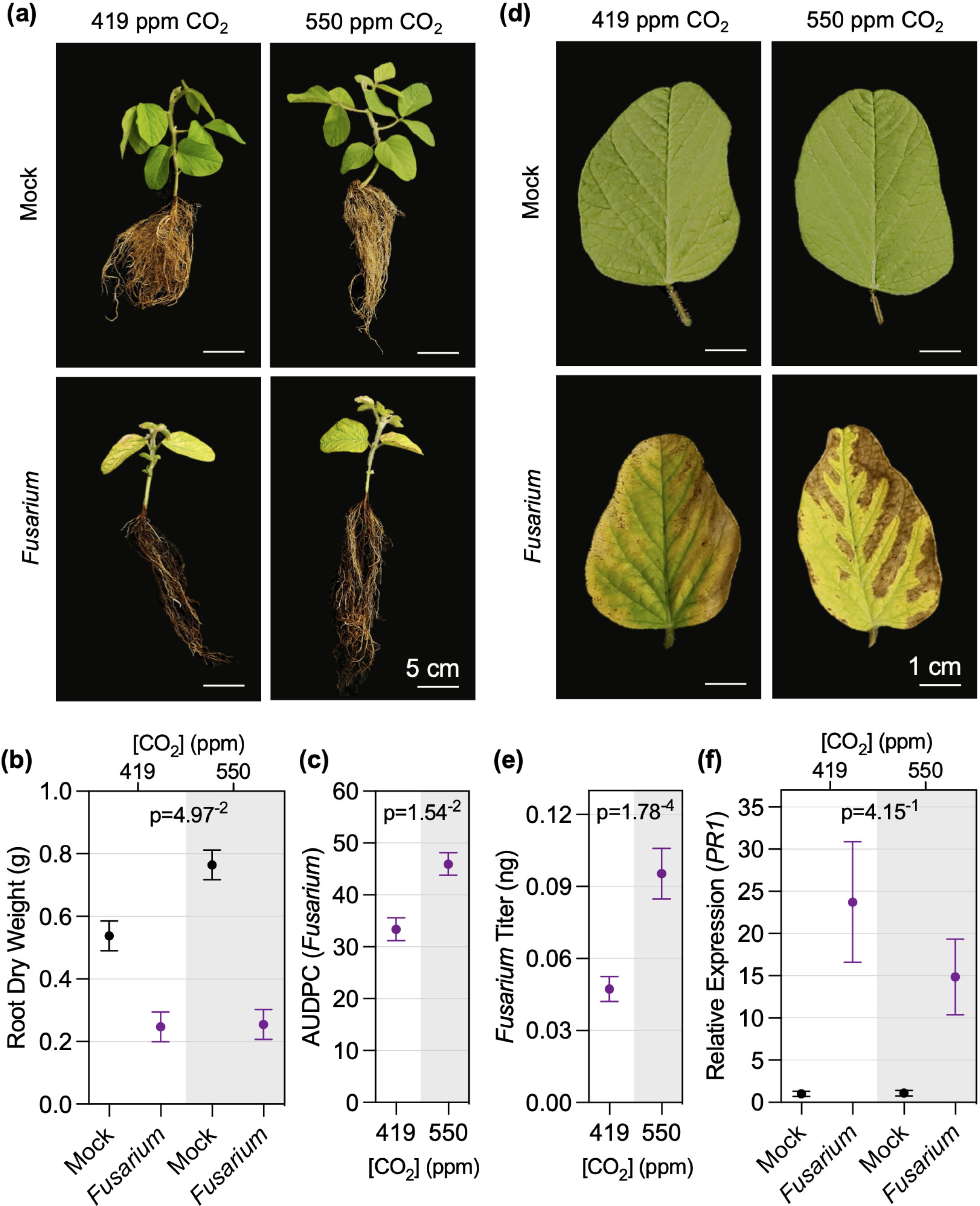
Young soybean plants growing in *e*CO_2_ develop more SDS symptoms and are more susceptible to *F. virguliforme*. **(a)** Representative photos of soybean plants at 35 days after planting (dap) that were mock treated or germinated in soil infested with *F. virguliforme* and **(b)** associated root dry weight. **(c)** SDS disease progression, assessed as area under the disease progress curve (AUDPC), and **(d)** close-up photos of SDS disease symptoms on unifoliate leaves at 35 dap. **(e)** *F. virguliforme* titer and **(f)** expression of *PR1* (SA marker gene) in soybean roots at 35 dap was assessed by RT-qPCR analysis relative to the *Skp1* housekeeping gene. The three experimental replicates were conducted simultaneously in independent CO_2_ controlled chambers using a replicated complete randomized design. Data was graphed as the mean across the three replicates with standard error bars. P-values were computed using F-tests for the main effect of CO_2_ on AUDPC and log-transformed titers, and interaction effect between CO_2_ and *Fusarium* treatment on root dry weight and log-transformed *PR1* gene expression from the LMM analysis.

Similar to *F. virguliforme*, *P. sylvaticum* caused root rot under both CO_2_ treatments (Figure 9a), and the root and shoot dry weights of infected plants were more dramatically decreased when compared to the mock treatment in *e*CO_2_ relative to *a*CO_2_ (Figure 9b, S11). Hyphal growth assays indicated that *P. sylvaticum* grew more quickly in *e*CO_2_ than in *a*CO_2_ (Figure S12). However, disease index ratings were similar between infected plants grown under both [CO_2_] (Figure 9c), which was consistent with the similar *P. sylvaticum* titers observed in roots at 14 dap (Figure 9d). Additionally, no difference was observed in *KTI1* expression (Figure 9e), and while the p-value for *PR1* expression was below 0.05, we do not consider it to be significant because of the similar levels of expression in *Pythium*-infected tissue (Figure S13) at 14 dap between the two [CO_2_]. Together, our results suggest that *e*CO_2_ could have a minor effect on *in vitro* growth of *P. sylvaticum*; however, we could not demonstrate that this led to the pathogen being more aggressive in the roots. It is also interesting that while susceptibility to *P. sylvaticum* appears to be unchanged there is a greater loss in root and shoot biomass in plants grown in *e*CO_2_.

**Figure 9.**
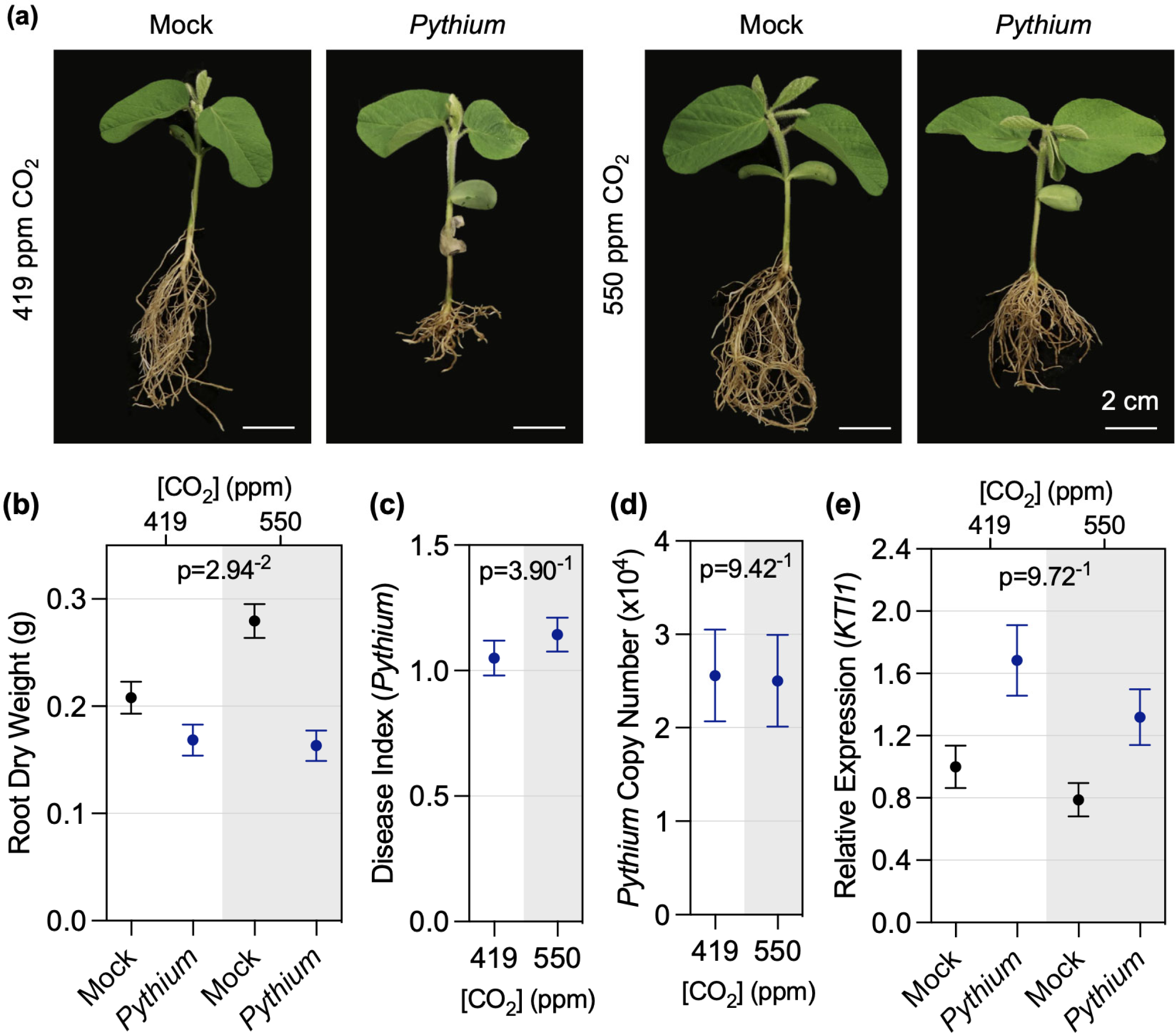
Effects of *e*CO_2_ on *P. sylvaticum* infection in soybean. **(a)** Representative photos of soybean plants at 21 days after planting (dap) that were mock treated or germinated in soil infested with *P. sylvaticum* and **(b)** associated root dry weight. **(c)** *P. sylvaticum* disease index measured 14 dap and **(d)** *P. sylvaticum* copy number in roots determined by RT-qPCR analysis at 21 dap. **(e)** *KTI1* (JA marker gene) expression in response to mock treatment or *P. sylvaticum* infection assessed by RT-qPCR analysis. Gene expression was assessed relative to the *Skp1* housekeeping gene. The three experimental replicates were conducted simultaneously in independent CO_2_ controlled chambers using a replicated complete randomized design. Data was graphed as the mean across the three replicates with standard error bars. P-values were computed using F-tests for the main effect of CO_2_ on AUDPC and *Pythium* copy number, and interaction effect between CO_2_ and *Pythium* treatment on root dry weight and log-transformed *KTI1* expression from the LMM analysis.

## Discussion

Climate change is impacting agriculture by affecting crop quality and yield (Pareek *et al*., 2020; Rezaei *et al*., 2023), and CO_2_ is considered to be the primary driver of climate change because of its abundance and longevity in the atmosphere. Given the central role of CO_2_ in plant biology, there are many ways it can influence plant physiology that can potentially impact interactions with pathogenic microbes (Bazinet *et al*., 2022). While some themes are beginning to emerge, it remains challenging to predict the response of a given plant to a given pathogen under *e*CO_2_. Here, we selected a diverse panel of phytopathogens to assess if and how soybean-microbe interactions are impacted by *e*CO_2_. Our results demonstrate that the CO_2_ levels expected within the next three decades (Jaggard *et al*., 2010) have significant impacts on soybean immunity. The effects of *e*CO_2_ on disease varied across the different pathogen types with enhanced resistance to bacterial blight caused by *Psg*, increased susceptibility to BPMV, SMV, and *F. virguliforme*, and little apparent effect on susceptibility to *P. sylvaticum*.

Extensive research from SoyFACE experiments has demonstrated that under favorable environmental conditions, *e*CO_2_ (550 ppm CO_2_) has the potential to benefit soybean growth and yield (Long *et al*., 2006; Ainsworth & Long, 2021). To determine if our *e*CO_2_ experiments in controlled environment chambers could approximate the effects on soybean growth and physiology that have been observed in SoyFACE experiments, we conducted baseline physiological and gene expression analyses. In line with field results, we observed that leaf photosynthetic rate and plant biomass increased, whereas stomatal conductance, density, and aperture decreased (Figure 1). Interestingly, we found that expression of some photosynthesis-related genes, such as *Rubisco small subunit 1B* (*RbcS1B*), decreased under eCO_2_, which seems contradictory to the enhanced photosynthesis. However, others have shown that mRNA expression of *RbcS* and other photosynthesis-related genes can be down-regulated as non-structural sugars accumulate and plants begin to acclimate to *e*CO_2_ (Cheng *et al*., 1998; Thompson *et al*., 2017). The combined results from physiology and baseline gene expression analyses indicated that the soybean plants were performing as expected in *e*CO_2_ under our growth conditions and support the idea that their responses to pathogens would be authentic.

Although the impacts of *e*CO_2_ on soybean physiology are well-established, its effects on disease are limited to one study that investigated the incidence and severity of three naturally occurring diseases caused by the fungal pathogens *F. virguliforme* (SDS) and *Septoria glycines* (brown spot) and the oomycete pathogen *Peronospora manshurica* (downy mildew) over three years in SoyFACE (Eastburn *et al*., 2010). *e*CO_2_ did not affect SDS incidence or severity, but the severity of brown spot disease increased, while the severity of downy mildew decreased. The incidence of brown spot and downy mildew were not affected. The results from this study are consistent with the idea that diseases caused by necrotrophic pathogens may be typically enhanced under *e*CO_2_, whereas diseases caused by biotrophic and hemibiotrophic pathogens may be suppressed in C3 plants (Bazinet *et al*., 2022). This study showed that future atmospheric CO_2_ levels have the potential to positively or negatively impact soybean diseases that occur in the field, but it was limited by the pathogens that were present, variable environmental conditions, and unknown levels of inoculum. Our goal was to extend these findings under controlled environmental conditions to determine how *e*CO_2_ differentially affects responses to distinct types of pathogens and to establish a foundation to understand the complex and dynamic signaling events that influence soybean-pathogen interactions in response to *e*CO_2_.

The effects of [CO_2_] on bacterial infection have not been investigated previously in soybean, however, several independent studies have indicated that for the hemibiotrophic pathogen *Pto*DC3000, *e*CO_2_ enhances resistance in tomato (Li *et al*., 2015; Zhang *et al*., 2015; Hu *et al*., 2021) and mostly increases susceptibility in Arabidopsis (Zhou *et al*., 2017, 2019). Arabidopsis does not become more susceptible to all pathogens, as it is more resistant to infection with the necrotrophic pathogen *B. cinerea* at *e*CO_2_ (Zhou *et al*., 2019). In our work, we observed enhanced resistance in compatible interactions between *Psg* and soybean under *e*CO_2_, and to a lesser extent, during incompatible interactions with *Pto*DC3000 and *Pto*DC3000 *hrcC^-^*(Figure 3A). Taken together, it is evident that bacterial immunity is altered under *e*CO_2_ in C3 plants in a pathosystem-specific manner. Notably, changes in environmental conditions, nutrient availability, and other growth conditions have been shown to alter the direction of these responses (Bazinet *et al*., 2022).

Phytohormones and stomatal immunity play pivotal roles in altering defense responses to bacteria under *e*CO_2_ in Arabidopsis and tomato. For example, in tomato, enhanced resistance to *Pto*DC3000 was associated with *e*CO_2_-induced stomatal closure (Li *et al*., 2015) and higher SA biosynthesis (Li *et al*., 2015; Zhang *et al*., 2015). In Arabidopsis, increased susceptibility to *Pto*DC3000 was associated with reduced ABA levels, causing changes in stomatal dynamics favoring bacterial infection (Zhou *et al*., 2017), and shifts in SA/JA antagonism toward JA-mediated immunity against necrotrophic pathogens (Zhou *et al*., 2019). In response to *Psg* infection, we observed a 1.7-fold increase in SA accumulation and no notable change in JA levels (Figure S4). However, this slight increase in SA seems unlikely to provide the level of resistance against *Psg* observed under *e*CO_2_ based on previous work showing that free SA was increased by 7-10 fold in soybean plants exhibiting SA-dependent, constitutive defense responses (Liu *et al*., 2011). We hypothesize that enhanced bacterial resistance is likely conferred by a combination of physiological, metabolic, and molecular responses that together limit bacterial infection under *e*CO_2_. Indeed, we observed an upregulation of MAMP-triggered immune responses in soybeans at *e*CO_2_ including flg22-induced oxidative species production (Figure 4b) and flg22-induced MAPK activation (Figure 4c). We additionally observed reduced stomatal density (Figure 1c) and a constitutive decrease in stomatal aperture (Figure 1d), which is likely to limit the entry of bacterial species.

Our transcriptomic analysis also indicates that *e*CO_2_ has direct impacts on soybean defense. *e*CO_2_ caused an over-representation of DEGs associated with defense responses (Figure 2, Table S5), even in the absence of pathogen challenge. Most notable was altered expression of genes associated with JA signaling and response to JA, including decreased expression of *Jasmonate Zim Domain* (*JAZ*)-encoding genes, which repress jasmonate signaling, and *MYC2* orthologs, known as the master regulators of jasmonate signaling (Johnson *et al*., 2023). Although we did not observe higher baseline JA levels under *e*CO_2_ in our metabolomics analysis (Figure S4), down-regulation of these genes suggests that JA-mediated responses may be altered in soybean plants growing in *e*CO_2_. This finding is consistent with prior work demonstrating that various insects prefer to feed on and cause more damage to plants grown in *e*CO_2_ (DeLucia *et al*., 2008; Johnson *et al*., 2020), which has been attributed at least in part to compromised JA-mediated defenses (Zavala *et al*., 2008). We also found no evidence for constitutive up-regulation of SA-mediated defenses under *e*CO_2_, because there were no pathogenesis-related genes among our DEGs and SA levels were not different from plants grown in *a*CO_2_. However, we did find that some genes that positively regulate basal defense responses were upregulated, suggesting that some immune responses might be primed in soybean under these conditions. The effects of *e*CO_2_ on DEGs was even more pronounced following *Psg* infection (Figure 5; Table S8). Similar DEGs were identified at *a*CO_2_ and *e*CO_2_ in response to *Psg* infection, however the amplitude of these responses were significantly more up- or down-regulated, depending on the gene, in response to *e*CO_2_ (Table S8). This is in line with the higher *Psg*- and flg22-elicited defense gene expression and basal defense responses under *e*CO_2_.

The effects of *e*CO_2_ on virus susceptibility has been studied in various other C3 plants, most of which have reported increased resistance to viral pathogens associated with a constitutive overproduction of SA (Fu *et al*., 2010; Trębicki *et al*., 2016) or upregulation of RNA silencing genes that are critical for viral defense (Guo *et al*., 2020). As discussed above, we did not observe a constitutive stimulation of SA levels or SA-based defenses in our experimental system, and interestingly, we found that expression of two genes potentially involved in anti-viral RNA silencing was decreased (Figure 7h,i). In line with our molecular analyses, soybean plants were more susceptible to BPMV and SMV in *e*CO_2_. Increased susceptibility could be partially explained by reduced induction of SA-based defenses, as indicated by reduced *PR1* gene expression in BPMV- and SMV-treated plants in *e*CO_2_ (Figure 7f). It is also possible that changes in RNA silencing components are responsible for increased susceptibility, as we observed a reduction in both *AGO1* (Figure 7h) and *DCL2* (Figure 7i) expression in virus-infected soybean at *e*CO_2_. Our results suggest that both SA-mediated defense and RNA silencing are compromised in response to BPMV and SMV infection in soybean. An interesting avenue to pursue in the future would be to test if there is a link between the apparent down-regulation of SA-mediated defenses and anti-viral RNA silencing.

Many viruses, including BPMV and SMV, are transmitted by an insect vector in nature. Importantly, many of the studies indicating that plants are less susceptible to viruses under *e*CO_2_ have used aphids as a vector for viral delivery. Thus, we do not know if the presence of bean leaf beetle (BPMV) or aphid (SMV) may influence the outcome of the interaction. Furthermore, the behavior of insect vectors or other factors may be influenced by *e*CO_2_ in ways that could affect dissemination of the viruses. For example, the larger canopy size associated with increased photosynthetic rates in soybean are expected to cause a higher dispersion rate, increasing disease incidence and severity in the field (Amari *et al*., 2021), pointing to viruses as potential pathogens of concern under predicted future climatic conditions. It will be interesting to study the tri-trophic interactions in the future from mechanistic and epidemiological perspectives, and perhaps including other climate change-associated gasses such as ozone, which retards SMV infection in soybean (Bilgin *et al*., 2008). *e*CO_2_ has been shown to have various effects on diseases caused by fungi and oomycetes (filamentous pathogens). Fungal pathogens are of particular concern under predicted future climatic conditions as it will allow their dispersal into more Northern latitudes as a consequence of higher temperatures and predicted changes in annual precipitation and humidity levels (Nnadi & Carter, 2021). While various studies have investigated the effect of *e*CO_2_ on filamentous pathogens (Smith & Luna, 2023), relatively few have examined soil-borne pathogens. Elevated CO_2_ (800 ppm) had no effect on infections caused by *Rhizoctonia solani* or *F. oxysporum* in Arabidopsis plants (Zhou *et al*., 2019). Tomato plants grown in 700 ppm CO_2_ were more tolerant to root rot caused by *Phytophthora infestans* (Jwa & Walling, 2001). Growth of *P. infestans* was not directly affected by [CO_2_] and the increased resistance of tomato plants could not be associated with changes in expression of marker genes associated with SA-, ABA, or JA-based defenses. In FACE experiments involving natural infections, *e*CO_2_ led to higher incidence of sheath blight (*Rhizoctonia solani*) on rice (Kobayashi *et al*., 2006), but SDS (*F. virguliforme*) in soybean was not affected (Eastburn *et al*., 2010). Interestingly, the increased incidence of rice sheath blight was only observed in plots that also received input of high nitrogen fertilizer. These previous studies show that responses of plants to filamentous soil-borne pathogens can be affected by *e*CO_2_, but they can be highly influenced by environmental factors and do not necessarily correlate with the expected defense-related signaling events.

Our data indicate that soybean plants are more susceptible to *F. virguliforme* under *e*CO_2_ but had no notable change in *P. sylvaticum* infection. Surprisingly, we observed a greater loss in root and shoot dry weight in response to *P. sylvaticum* even though visual disease scores were similar, and similar quantities of *P. sylvaticum* DNA were observed in the roots of plants grown at *e*CO_2_ and *a*CO_2_. We only observed a modest reduction in *PR1* expression following *F. virguliforme* treatment under *e*CO_2_, and there was no change in expression of the *KTI1* JA marker gene. These gene expression results are similar to what has previously been observed in tomato with increased susceptibility showing no correlation with concomitant changes in expression of defense pathway marker genes (Jwa & Walling, 2001). Various other factors have been linked to *F. virguliforme* resistance, which we did not assess in this study, including the production of phytoalexins and changes in root exudates, both of which can be affected by *e*CO_2_ (Vaughan *et al*., 2014; Usyskin-Tonne *et al*., 2020). Experiments at FACE facilities have demonstrated dramatic changes in the soil microbial rhizosphere and endosphere (Rosado-Porto *et al*., 2022; Gao *et al*., 2022), which could also impact the composition of phytopathogenic fungi in the surrounding soil and cause changes in soybean-pathogen interactions. One could also expect that changes in root architecture and growth could benefit or curtail infection by soil pathogens. Our data indicate that disease losses due to soil-borne filamentous pathogens can be exacerbated in soybean grown in *e*CO_2_, and there is much more to learn about the underlying mechanisms.

In conclusion, we have demonstrated that soybean basal defense responses and gene expression are altered under *e*CO_2_, differentially regulating susceptibility to bacterial, viral, fungal, and oomycete pathogens. Comprehensive analysis of bacterially-elicited responses has given us a strong foundation to investigate which genes are regulating defense to bacteria under *e*CO_2_ and how the corresponding signaling pathways are regulated. In our experiments, we assessed the effect of [CO_2_] on pathogen growth, however, we did not investigate how pathogen virulence or evolution affects soybean-pathogen interactions under these conditions. In our study, we focused on the impact of one factor, *e*CO_2_, on soybean physiology and disease susceptibility. However, *e*CO_2_ is not the only environmental factor to consider when predicting the outcome of future climatic conditions on plant health. Increases in temperature, changes in water and nutrient availability, soil pH, photoperiod, and severe weather events are also expected to affect disease susceptibility (Saijo & Loo, 2020). For example, higher temperatures have almost uniformly been associated with higher disease susceptibility to bacterial pathogens in Arabidopsis (Cohen & Leach, 2020). Understanding how combined stresses, in the context of future atmospheric CO_2_ levels that are likely to occur, affect defense signaling and disease development is a critical question that needs to be investigated further to develop new strategies to mitigate crop losses due to threats imposed by climate change.

## Acknowledgments

We acknowledge the ISU DNA Facility, Office of Biotechnology, Iowa State University, Ames IA for providing analytical instrumentation and technical expertise. We acknowledge the Iowa State University W.M. Keck Metabolomics Research Laboratory (RRID:SCR_017911) for providing analytical instrumentation. We thank Dr. Gwyn Beattie (ISU) for providing us with the *Pto*DC3000 *hrcC-* strain, Dr. Leanor Leandro and Vijitha K. Silva (ISU) for SDS inoculum and inoculation protocols, and Dr. Alison E. Robertson and Clarice L. Schmidt (ISU) for *P. sylvaticum* inoculum and guidance with infection experiments. This work was supported in part by the Iowa Soybean Research Center, the ISU Plant Sciences Institute, USDA NIFA Hatch Project 4308, the ISU College of Agriculture and Life Sciences and by USDA-ARS projects 5030-21220-007-000D (Leveraging Crop Genetic Diversity and Genomics to Improve Biotic and Abiotic Stress Tolerance in Soybean) and 0500-00093-001-00-D (SCINet). Ekkachai Khwanbua is supported by an Anandamahidol fellowship from the government of Thailand. Mention of trade names or commercial products in this publication is solely for the purpose of providing specific information and does not imply recommendation or endorsement by the USDA. USDA-ARS is an equal opportunity employer and provider.

## Author Contributions

EK, MB, ASC, and KH performed the experiments. MB, EK, and SAW conceived of the experiments. MWB performed the metabolomics analysis. YQ and PL performed statistical analysis using the linear mixed models. MAG performed the QuantSeq data analysis. MB, EK, MAG, and SAW wrote the manuscript with contributions from all authors.

## Data Availability

The RNA-seq reads for QuantSeq dataset 1 and dataset 2 were deposited in NCBI under BioProject PRJNA1017882 and PRJNA1017884, respectively. The authors declare that any additional supporting data for this work can be found in the manuscript, its supplementary materials, or obtained directly from the corresponding author upon request.

