## Supplemental Figures 1 - 13 for "Elevated CO_2_ alters soybean physiology and defense responses, and has disparate effects on susceptibility to diverse microbial pathogens"

### Supporting Information

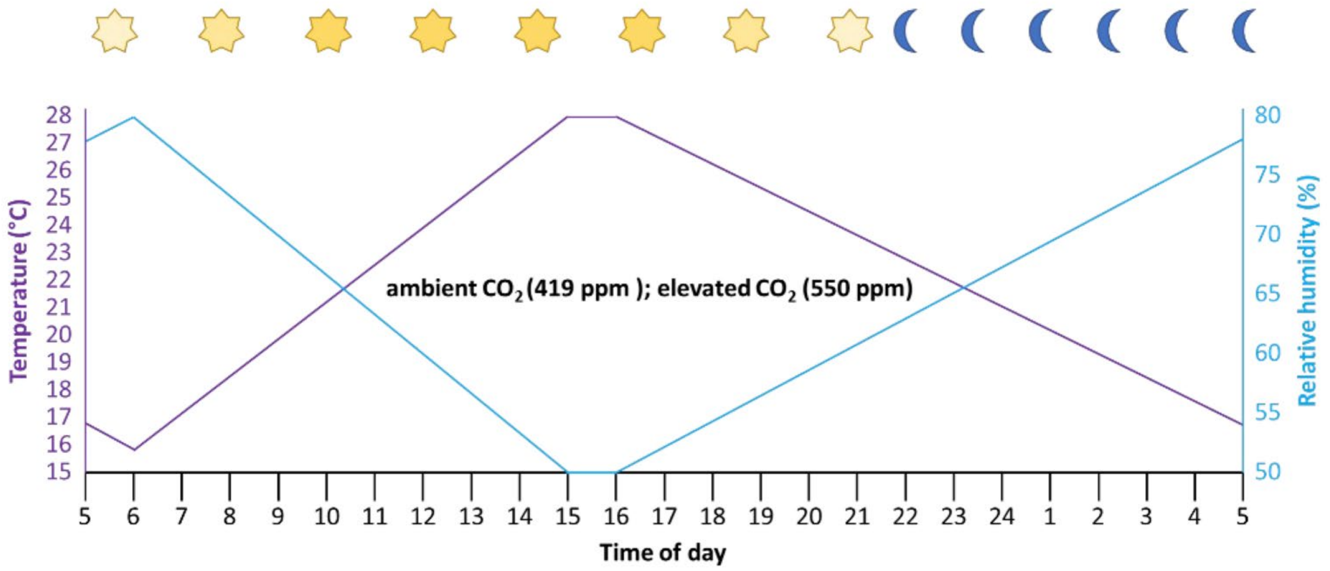


**Figure S1. Schematic representation of growth chamber conditions.** Schematic representation of Enviratron growth chamber conditions. All chambers were set to the same conditions, except that $CO_2$ was 419 ppm in chambers set to ambient (*a*$CO_2$) conditions and $CO_2$ was 550 ppm in elevated (*e*$CO_2$) conditions. Lights were on at 5:38 am (05:38) and off at 8:52 pm (22:52) (sunrise and sunset times represent June 15 in central Iowa), minimum temperature was 15.6 °C at 6 am (06:00) with ramping to 27.6 °C maximum temperature from 3:00 – 4:00 pm (15:00 – 16:00). The relative humidity was ramped between 80% at 6:00 am (0:600) to the minimum of 50% at 3:00 – 4:00 pm (15:00-16:00).

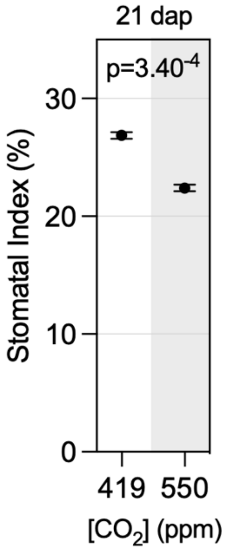


**Figure S2. Stomatal index on the abaxial leaf surface of soybean.** Stomatal index measurements were performed as described in materials and methods. Three replicates were carried out simultaneously in independent $CO_2$ control chambers, using 5 plants per treatment for each $CO_2$ concentration. Stomata were counted in three randomly selected fields of view per leaf. Data points represent the mean value across the three replicates, and error bars represent the standard error. P-value was computed based on an F-test for the main effect of $CO_2$ in linear mixed effect model analysis.

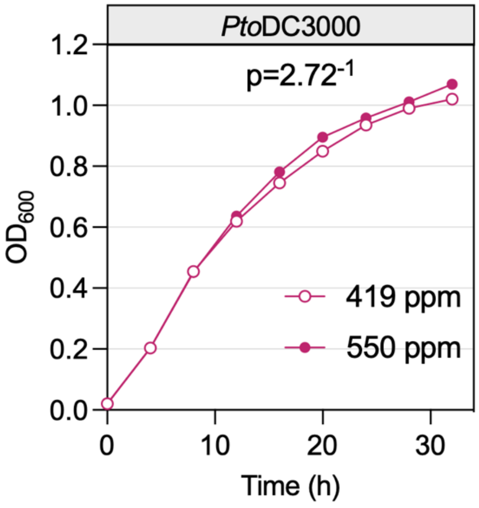


**Figure S3. Effect of elevated $CO_2$ on growth rate of *Pseudomonas syringae* pv. *tomato* DC3000 (*Pst*DC3000).** Bacterial cultures were grown in LB broth without antibiotics for 32 h and $OD_{600}$ was measured every four hours using a portable spectrophotometer. Two experimental replicates were conducted simultaneously, for each $CO_2$ concentration, in independent $CO_2$ chambers. Data points represent the average $OD_{600}$ between the two chambers. P-value was computed based on an F-test for the interaction of $CO_2$ and time in linear mixed effect model analysis using data after 4 hours due to unequal variance.

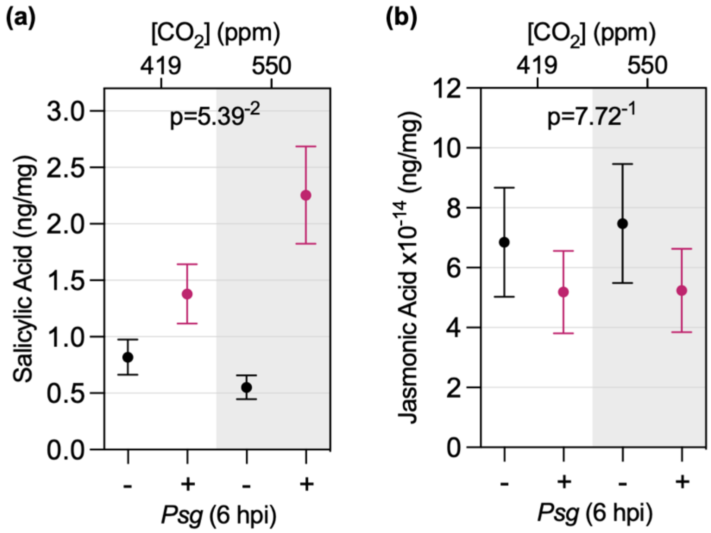


**Figure S4. Accumulation of SA and JA at 6 h post-inoculation with *Pseudomonas syringae* pv. *glycinea* (*Psg*) in *a*$CO_2$ and *e*$CO_2$. (a)** SA and **(b)** JA levels were quantified using non-targeted and targeted metabolomics analysis, respectively, as described in methods and materials. Experiments were conducted independently, three times, using 8 plants per treatment for each $CO_2$ concentration. Data points represent the mean value across the three replicates and error bars represent the standard error. P-value was computed based on an F-test for the interaction of $CO_2$ and *Psg* treatment in linear mixed effect model analysis on log-transformed data.

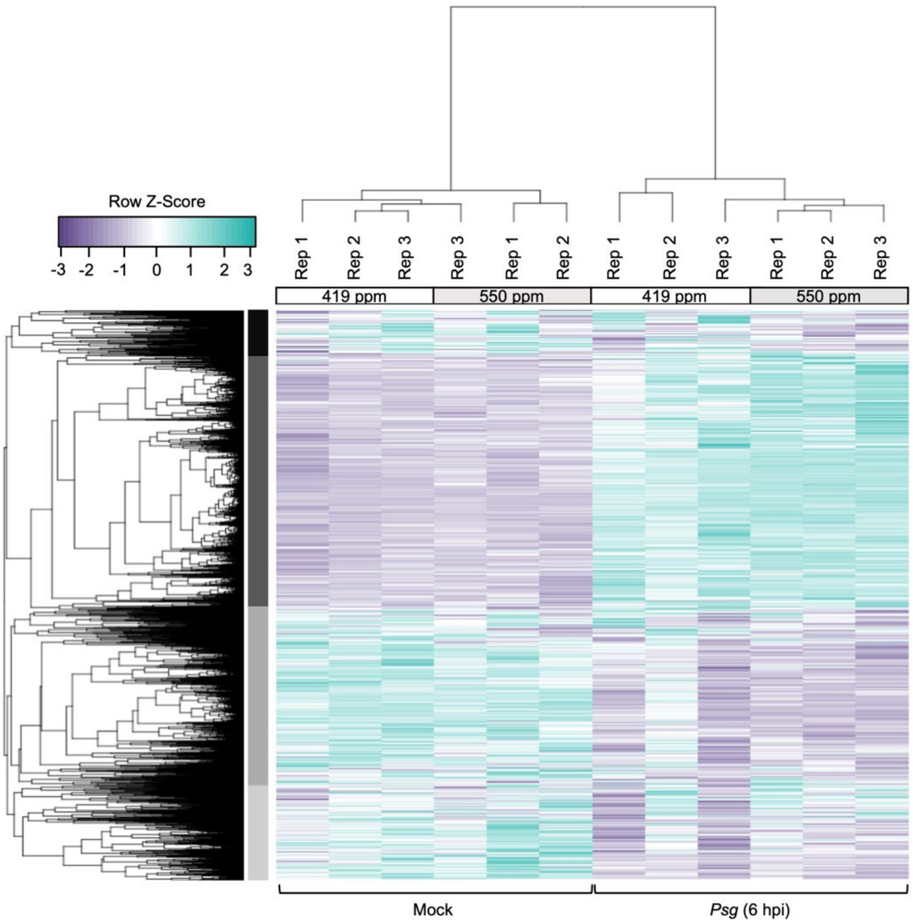


**Figure S5. Hierarchical clustering of 21,822 DEGs (FDR < 0.01) responding to *Pseudomonas syringae* pv. *glycinea* (*Psg*) at 6 h post-inoculation in plants grown in *e*$CO_2$ (550 ppm) or *a*$CO_2$ (419 ppm) conditions.** The full list of DEGs is provided in Table S7. Samples for QuantSeq analysis were taken from the unifoliate leaves of 14-day old plants at 6 h post-inoculation with mock treatment or *Psg*. The three independent replicates were conducted using eight plants per $CO_2$ treatment. Rep indicates independent biological replicates. Row Z-scores were used for hierarchical clustering of DEGs, based on expression across samples and replicates. Purple indicates expression values below the row mean and teal indicates expression values above the row mean. Four different expression clusters were identified. These are indicated by differently shaded gray boxes to the left of the heat map.

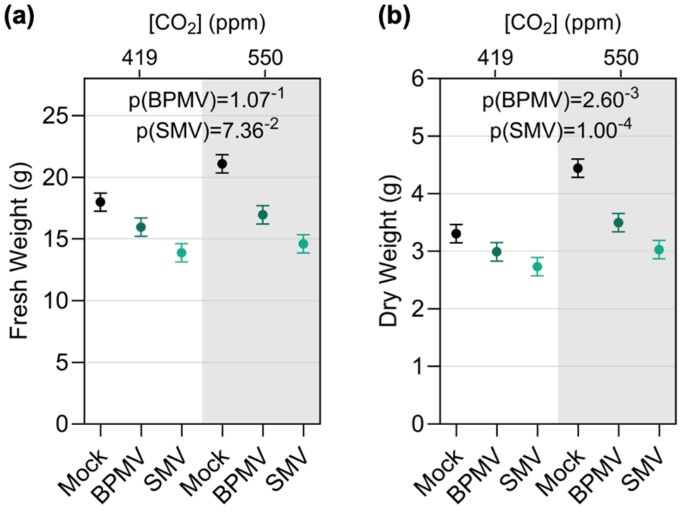


**Figure S6. Effect of elevated $CO_2$ on the shoot fresh weight and shoot dry weight of soybean plants infected with bean pod mottle virus (BPMV) and soybean mosaic virus (SMV). (a)** Fresh weight and **(b)** dry weight of soybean plants at 21 days post-inoculation with BPMV, SMV, or mock treatment. Three experiments were conducted simultaneously in independent $CO_2$ control chambers, using 8 plants per treatment for each $CO_2$ concentration according to a replicated complete design. Data points represent the mean value across the three replicates and error bars represent the standard error. P-values were computed based on t-tests for comparing the difference between virus and Mock under $eCO_2$ with that under $aCO_2$ in linear mixed effect model analysis.

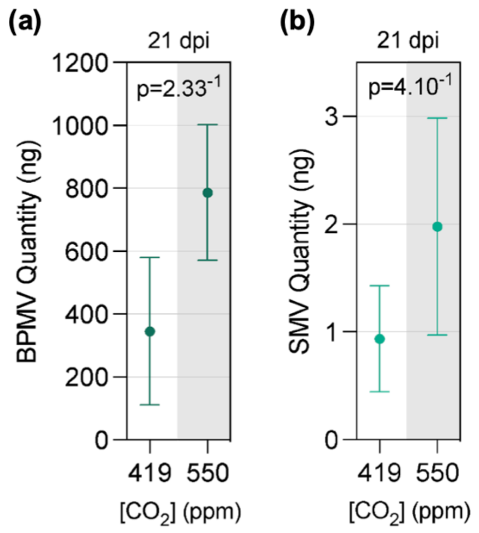


**Figure S7. Bean pod mottle virus (BPMV) and soybean mosaic virus (SMV) accumulation at 21 days post-inoculation in soybean plants growing in *a*$CO_2$ and *e*$CO_2$. (a)** BPMV and **(b)** SMV quantity was determined by RT-qPCR using multiplexed probes for BPMV or SMV and *Skp1* as an internal reference control. Samples were taken from the newest fullest expanded leaf. Three experiments were conducted simultaneously in independent $CO_2$ chambers, using 8 plants per infection treatment for each $CO_2$ concentration according to a replicated complete block design. Data points and bars represent mean group values and standard error across the three replicates. P-value for BPMV was computed based on an F-test for the main effect of $CO_2$ in linear mixed effect model analysis. P-value for SMV was computed based on an F-test for the main effect of $CO_2$ in linear mixed effect model analysis on log-transformed data.

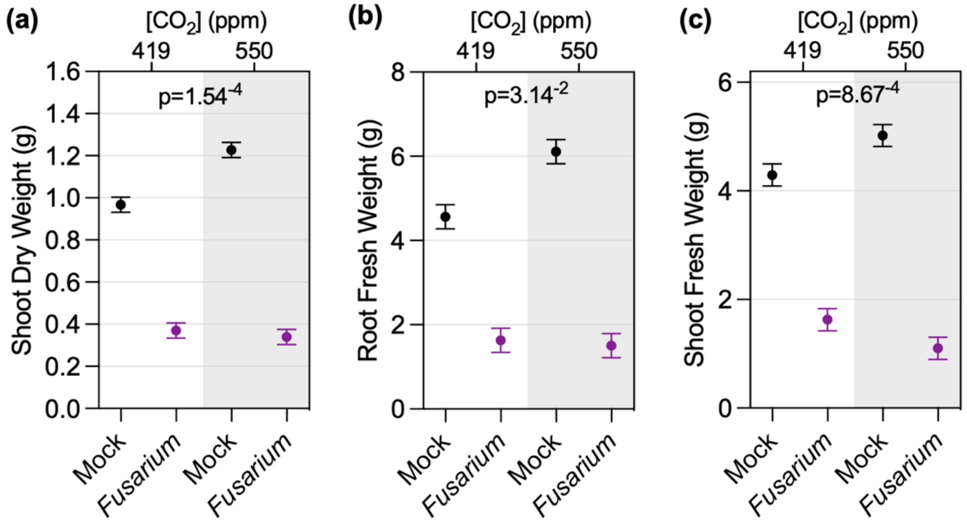


**Figure S8. Effects of *e*$CO_2$ on root and shoot biomass of soybean plants infected with *Fusarium virguliforme*.** **(a)** Shoot dry weight, **(b)** root fresh weight and **(c)** shoot fresh weight at 35 days after germination in soil infested with *F. virguliforme* or mock infested. The three replicate experiments were conducted simultaneously in independent $CO_2$ control chambers using 6 plants per treatment for each $CO_2$ condition according to a replicated complete block design. Data points represent mean values with standard error across the three replicates. P-values were computed based on F-tests for the interaction effect between $CO_2$ and *Fusarium* treatment in linear mixed effect model analysis.

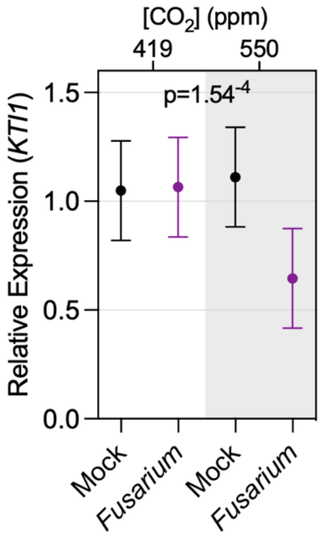


**Figure S9. Expression of the JA marker gene, *KTI1*, in roots of plants infected with *Fusarium virguliforme* in *a*$CO_2$ and *e*$CO_2$.** RT-qPCR analysis was conducted on RNA extracted from root tissue using gene-specific primers for the JA marker gene *KTI1* with *Skp1* as an internal reference control. Experiments were conducted using 6 plants per treatment for each $CO_2$ concentration. Triplicate experiments were conducted simultaneously in independent $CO_2$ chambers. Data points and bars represent mean values and standard error across the three replicates. P-value was computed based on an F-test for the interaction effect between $CO_2$ and *Fusarium* treatment in linear mixed effect model analysis.

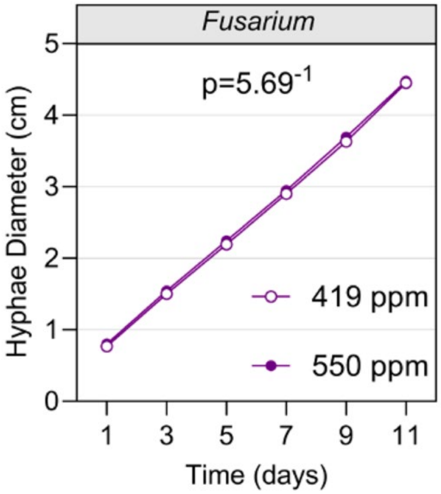


**Figure S10. $CO_2$ concentration does not impact *in vitro* growth of *Fusarium virguliforme*.** Hyphae diameter was measured every 2 days over 11 days. Three experiments were conducted simultaneously using 10 plates per experiment, for each $CO_2$ treatment. Data points represent mean values across three replicates conducted simultaneously in independent $CO_2$ control chambers. P-value was computed based on an F-test for the interaction effect between $CO_2$ and time in linear mixed effect model analysis.

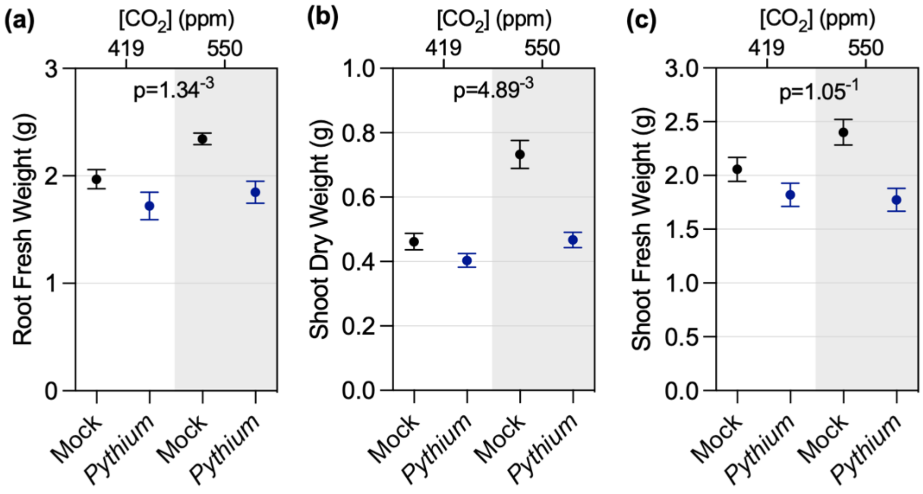


**Figure S11. Soybean root and shoot biomass is impacted by *e*$CO_2$ in *Pythium sylvaticum* infected plants (a)** Root fresh weight, **(b)** shoot dry weight and **(c)** shoot fresh weight in mock or *P. sylvaticum*-infected plants 21 days after inoculation. Experiments were conducted in triplicate using 6 plants per treatment, for each $CO_2$ condition. Triplicate experiments were conducted simultaneously in independent $CO_2$ controlled chambers. Data points represent mean values across the three replicates with standard error bars. P-values were computed based on F-tests for the interaction effect between of $CO_2$ and *Pythium* treatment in linear mixed effect model analysis. Fourth power and log transformation were applied for root fresh weight and shoot dry weight respectively due to unequal variance on the original scale.

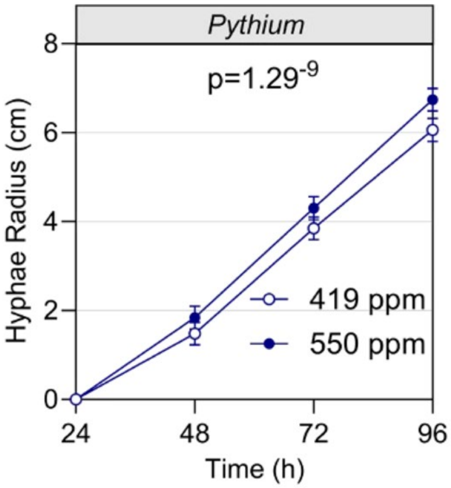


**Figure S12. *In vitro* growth of *Pythium sylvaticum* is accelerated under *e*$CO_2$.** Experiments were conducted using 10 plates per treatment for each $CO_2$ condition in triplicate. The three experiments were done simultaneously in independent $CO_2$ controlled chambers. Hyphae radius was measured every 4 h over 32 h. Data points and bars represent mean values and standard error across the three replicates. P-value was computed based on an F-test for the interaction effect between $CO_2$ and time in linear mixed effect model analysis.

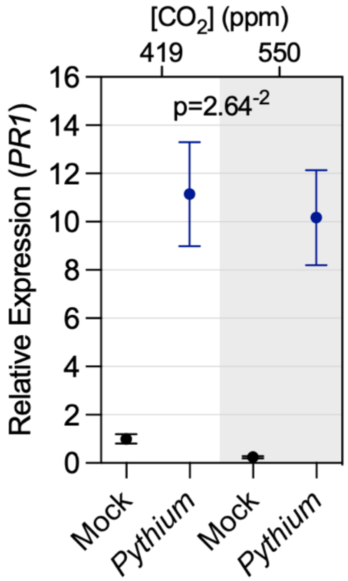


**Figure S13. Expression of the SA marker gene, *PR1*, in roots of plants infected with *Pythium sylvaticum* in *a*$CO_2$ and *e*$CO_2$.** RT-qPCR analysis was conducted using RNA extracted from soybean roots 14 days after inoculation using gene specific primers for the SA marker gene *PR1* and *Skp1* was an internal reference control. Experiments were conducted simultaneously in three independent $CO_2$ controlled chambers using 6 plants per treatment for each $CO_2$ concentration according to a replicated complete block design. Data points represent mean values across the three replicates with standard error. P-value was computed based on an F-test for the interaction effect between $CO_2$ and *Pythium* treatment in linear mixed effect model analysis on log-transformed data.
